# Bi-directional flow of the funny current (I_f_) during the pacemaking cycle in murine sinoatrial node myocytes

**DOI:** 10.1101/2021.03.10.434820

**Authors:** Colin H. Peters, Pin W. Liu, Stefano Morotti, Stephanie C. Gantz, Eleonora Grandi, Bruce P. Bean, Catherine Proenza

**Author notes:** Corresponding Author. Catherine Proenza, PhD Department of Physiology & Biophysics University of Colorado – Anschutz Medical Campus 12800 E. 19th Ave, MS 8307 Aurora, CO 80045.

## Abstract

Sinoatrial node myocytes (SAMs) act as cardiac pacemaker cells by firing spontaneous action potentials (APs) that initiate each heartbeat. The funny current, I_f_, is critical for the generation of these spontaneous APs; however, its precise role during the pacemaking cycle remains unresolved. We used the AP-clamp technique to quantify I_f_ during the cardiac cycle in mouse SAMs. We found that I_f_ is persistently active throughout the sinoatrial AP, with surprisingly little voltage-dependent gating. As a consequence, it carries both inward and outward current around its reversal potential of -30 mV. Despite operating at only 2-5% of its maximal conductance, I_f_ carries a substantial fraction of both depolarizing and repolarizing net charge movement during the firing cycle. We also show that β-adrenergic receptor stimulation increases the percentage of net depolarizing charge moved by I_f_, consistent with a contribution of I_f_ to the fight-or-flight increase in heart rate. These properties were confirmed by heterologously-expressed HCN4 channels and by mathematical models of I_f_. Modelling further suggested that the slow activation and deactivation of the HCN4 isoform underlie the persistent activity of I_f_ during the sinoatrial AP. These results establish a new conceptual framework for the role of I_f_ in pacemaking, in which it operates at a very small fraction of maximal activation but nevertheless drives membrane potential oscillations in SAMs by providing substantial driving force in both inward and outward directions.

**Significance Statement:** Cardiac pacemaker cells trigger each heartbeat by virtue of spontaneous oscillations in their membrane voltage. Although the funny current (If) is critical for these oscillations and for setting heart rate, its precise role remains an enigma because it activates mostly outside of the physiological voltage range and quite slowly relative to the pacemaker cycle. Here we show that I_f_ is persistently active in pacemaker cells; once opened, the small fraction of ion channels that conduct I_f_ do not re-close. Consequently, I_f_ flows both inward and outward to help propel the voltage oscillations and it paradoxically conducts a large fraction of the net charge movement. These results establish a new conceptual framework for the role of I_f_ in driving cardiac pacemaking.

## Introduction

Each beat of the heart is initiated by spontaneous pacemaker activity that originates in the sinoatrial node (1). Sinoatrial node myocytes (SAMs) are specialized cardiomyocytes that function as pacemaker cells by generating spontaneous action potentials (APs). Sinoatrial APs differ considerably from the APs of working cardiomyocytes in the atrial and ventricular myocardium, reflecting the unique function and protein expression of SAMs. In addition to having a slower upstroke velocity and smaller amplitude, sinoatrial APs are characterized by a spontaneous depolarization during diastole that drives the membrane potential to threshold to trigger the next AP.

The funny current, I_f_, is a hallmark of SAMs and is among the many ionic currents that are known to contribute to the generation of spontaneous APs in SAMs (2). I_f_ was first described in the late 1970s as a hyperpolarization-activated inward current that is increased by β-adrenergic receptor stimulation (3). I_f_ is produced by hyperpolarization-activated cyclic nucleotide-sensitive (HCN) channels, which were first cloned in the late 1990s (4). HCN4 is the primary HCN channel isoform expressed in the sinoatrial node of all mammals (5, 6). I_f_ and HCN4 are known to be critical for pacemaking — pharmacological inhibition of I_f_ slows heart rate (7, 8) and mutations in HCN4 cause sinus arrhythmias in humans and animal models (9–11). However, there is still debate about the specific contribution of If to the sinoatrial AP, in part because the steady-state voltage-dependence and slow activation of the current seemingly preclude appreciable channel activity at physiological potentials during the dynamic pacemaker cycle (12, 13).

I_f_ is generally described as an inward current and its role in pacemaking is typically considered only in terms of the diastolic depolarization phase of the sinoatrial AP (9, 12, 14–16).

However, its slow kinetics (6, 17), the presence of a voltage-independent component to HCN channel current (18, 19), and permeability to both Na^+^ and K^+^ with a net reversal potential around -30 mV suggest the possibility that I_f_ may remain active throughout the duration of the sinoatrial AP, and that it may also conduct outward current. Indeed, in a review of eleven different computational models of I_f_, Verkerk and Wilders show that all models produce some outward current during the upstroke and repolarization phases of the AP (20), despite the large variations between models due to differences in experimental conditions of the underlying data. Thus, I_f_ may also contribute to the repolarization phase of the sinoatrial AP, an important functional role that is typically not assayed in experiments.

Uncertainty about the role of I_f_ in shaping the sinoatrial AP is driven by the difficulty of directly measuring the current during the firing cycle. A fundamental limitation to studying the contribution of any individual current in SAMs is that manipulation of a single current — for example, by blockers or genetic knockout — alters the AP morphology and the activation of other currents, which in turn feeds back to change the current of interest. This difficulty is further compounded by a large variability in sinoatrial AP waveforms and I_f_ amplitudes, even within cells isolated from the same animal (21–23). This is to say nothing of the large effects that physiological changes such as age can have on the AP (24, 25). And, finally, the measurement of I_f_ is limited by the lack of specific blockers that can be used to isolate I_f_ from other currents. Indeed, HCN channel blockers, such as ivabradine, ZD7288, zatebradine, cilobradine, and Cs^+^, have been shown to also inhibit voltage-gated Ca^2+^, K^+^, and Na^+^ channels (26–35).

To overcome these challenges and directly measure I_f_ during the sinoatrial AP, we developed a series of pharmacological blockers to minimize the off-target effects of ivabradine and then used the AP clamp technique to isolate I_f_ as the ivabradine-sensitive current flowing in response to the AP in acutely-isolated mouse SAMs. Our recordings revealed that I_f_ in SAMs is biphasic, flowing both inward and outward during the AP. The trajectory of I_f_ tracked that of the membrane potential remarkably closely, and current-voltage relationships indicated that there is little voltage-dependent gating of I_f_ during the sinoatrial AP. These results were confirmed by heterologously-expressed HCN4 channels and by mathematical models of If. Together the data suggest a new conceptual framework for the role of I_f_ in pacemaking, in which it operates at a very small fraction of maximal activation but nevertheless drives membrane potential oscillations in SAMs by providing substantial driving force in both inward and outward directions.

## Results

### Isolation of I_f_ as an ivabradine-sensitive current in the presence of Na^+^, Ca^2+^ and K^+^ channel blockers

In order to directly measure I_f_ during the AP in SAMs, we used the FDA-approved drug ivabradine as a blocker because it appears to have somewhat higher specificity compared to other I_f_ inhibitors (32, 36, 37). To assess the degree of block, acutely-isolated SAMs from mice were voltage-clamped in the whole-cell configuration at 35°C and I_f_ was assayed in response to hyperpolarizing voltage steps before and during extracellular perfusion of ivabradine. We found that 30 μM ivabradine was required to achieve ∼90% block of I_f_ within ∼120 s of perfusion, consistent with its IC_50_ of ∼3 μM and intracellular site of action (36, 38–40) (Fig 1A, B; Table S1).^1^ In current-clamp experiments, we found that 30 μM ivabradine reduced AP firing rate by ∼50% within 30 s of perfusion (Fig. S1; P < 0.0001) and completely stopped spontaneous APs within 120 s of perfusion in most (10 of 12) cells.

**Figure 1:**
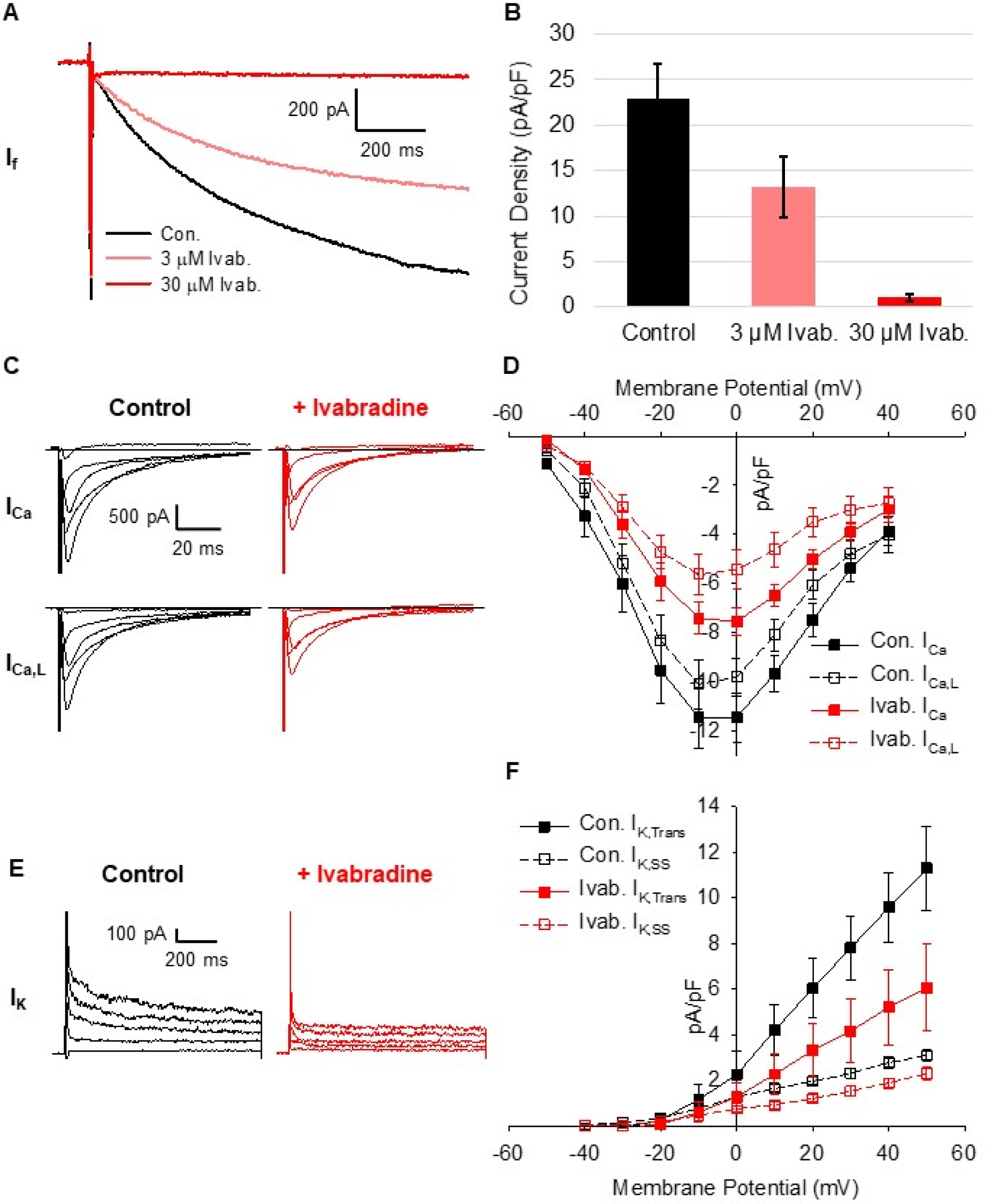
30 µM ivabradine blocks Ca^2+^ and K^+^ currents in SAMs. **A:** Representative whole-cell currents recorded from a murine SAM in response to 1 s hyperpolarizing voltage steps to -130 mV from a holding potential of -60 mV in a control Tyrode’s solution with 1 mM BaCl_2_ (*black*) and after perfusion of Tyrode’s containing 3 µM (*pink*) or 30 µM (*red*) ivabradine. **B:** Average (± SEM) current density of I_f_ in 1 mM BaCl_2_ (*black*) and after perfusion of 3 µM (*pink*) or 30 µM (*red*) ivabradine during pulses to -130 mV in murine SAMs. **C:** Representative total (*top*) and L-type (*bottom*) whole-cell Ca^2+^ currents recorded from sinoatrial myocytes in control (*black*) and after perfusion of 30 µM ivabradine (*red*). **D:** Mean (± SEM) total (*closed*) and L-type (*open*) Ca^2+^ current density in control and after perfusion of 30 µM ivabradine using the same colour scheme as **C**. **E:** Representative whole-cell K^+^ currents in a SAMs in control conditions (*black*) and after perfusion of 30 µM ivabradine (*red*). **F:** Average (± SEM) transient (*closed*) and steady-state (*open*) K^+^ current density in control conditions and after perfusion of 30 µM ivabradine using the same colour scheme as **E**. Horizontal lines indicate zero current. Number of replicates and details of statistical tests can be found in Table S1.

**Table 1:** Modelling parameters for equations 1-11.

| | 1 nM Isoproterenol<br>model | 1 μM Isoproterenol<br>model | Units |
| --- | --- | --- | --- |
| $G_f$ | 11.79 | | nS |
| $E_q$ | -35.4 | | mV |
| $E_{Na}$ | 70.49 | | mV |
| $E_K$ | -86.96 | | mV |
| $V_{1/2}$ | 103.18 | 88.53 | mV |
| $z$ | 11.6 | 9.15 | |
| $\tau_{f0}$ | 0.7 | 0.035 | ms |
| $k_f$ | 0.025 | 0.04 | $mV^{-1}$ |
| $\alpha_{f,1/2}$ | 340 | 330 | mV |
| $\beta_{f,1/2}$ | -160 | -170 | mV |
| $\tau_{s0}$ | 70 | 10 | ms |
| $k_s$ | 0.015 | 0.025 | $mV^{-1}$ |
| $\alpha_{s,1/2}$ | 300 | 285 | mV |
| $\beta_{s,1/2}$ | -200 | -215 | mV |
| $a$ | 0.015 | 0.003 | |
| $b$ | 0.025 | 0.03 | |
| $c$ | 300 | 330 | mV |
| $d$ | -200 | -170 | mV |

Where $E_{Na}$ and $E_K$ were calculated using the Nernst equations with 140 mM extracellular $Na^+$ , 10 mM intracellular $Na^+$ , 5.4 mM extracellular $K^+$ , and 140 mM intracellular $K^+$ .

Given previous reports of off-target actions of ivabradine (26, 34, 35), we next asked whether 30 μM ivabradine blocked an appreciable fraction of Ca^2+^ and K^+^ currents in SAMs under our recording conditions. Ca^2+^ currents were recorded in cesium-based solutions before and after perfusion of 30 μM ivabradine (Fig. 1C). Total whole-cell Ca^2+^ current was elicited using depolarizing voltage steps from a holding potential of -90 mV. L-type Ca^2+^ current was then measured in the same cells from a holding potential of -60 mV where T-type Ca^2+^ currents are mostly inactivated in SAMs (24, 41). We found that 30 μM ivabradine significantly reduced total and L-type Ca^2+^ currents, blocking 30-40% of peak current at 0 mV (Fig 1D; Table S1). T-type Ca^2+^ current was estimated in each cell by subtracting the L-type from the total Ca^2+^ current. Ivabradine had no significant effect on the calculated T-type current (Table S1). We found that 30 μM ivabradine also blocked a significant fraction of both the transient and steady-state K^+^ currents (∼46 and 32%, respectively, measured by a step to +40 mV from a holding voltage of - 50 mV; Fig. 1E, F; Table S1).

The above data indicate that ivabradine-sensitive currents in SAMs under physiological ionic conditions will include significant components of voltage-gated Ca^2+^ and K^+^ currents, particularly at more depolarized potentials. Thus, we next sought to define a series of blockers that could be used to pre-block Ca^2+^ and K^+^ currents without affecting I_f_ so that subsequent application of ivabradine could be used to isolate If. We measured Ca^2+^ currents before and after perfusion of 3 μM of the Ca^2+^ channel blocker isradipine (Fig. 2A). As expected, 3 μM isradipine blocked most (∼90%) of the peak total and L-type Ca^2+^ currents (Fig. 2B; Table S2). K^+^ currents were effectively inhibited by a cocktail of K^+^ channel blockers (10 mM TEA, 1 mM barium, and 1 µM E4031) followed by the same solution also containing 30 μM ivabradine (Fig. 2C). When ivabradine was applied after the K^+^ channel blockers, it had almost no effect on the peak transient K^+^ current and produced a small (but not significant) reduction of the steady-state K^+^ current that remained in the cocktail of K^+^ channel blockers (Fig. 2D; Table S2).

**Figure 2:**
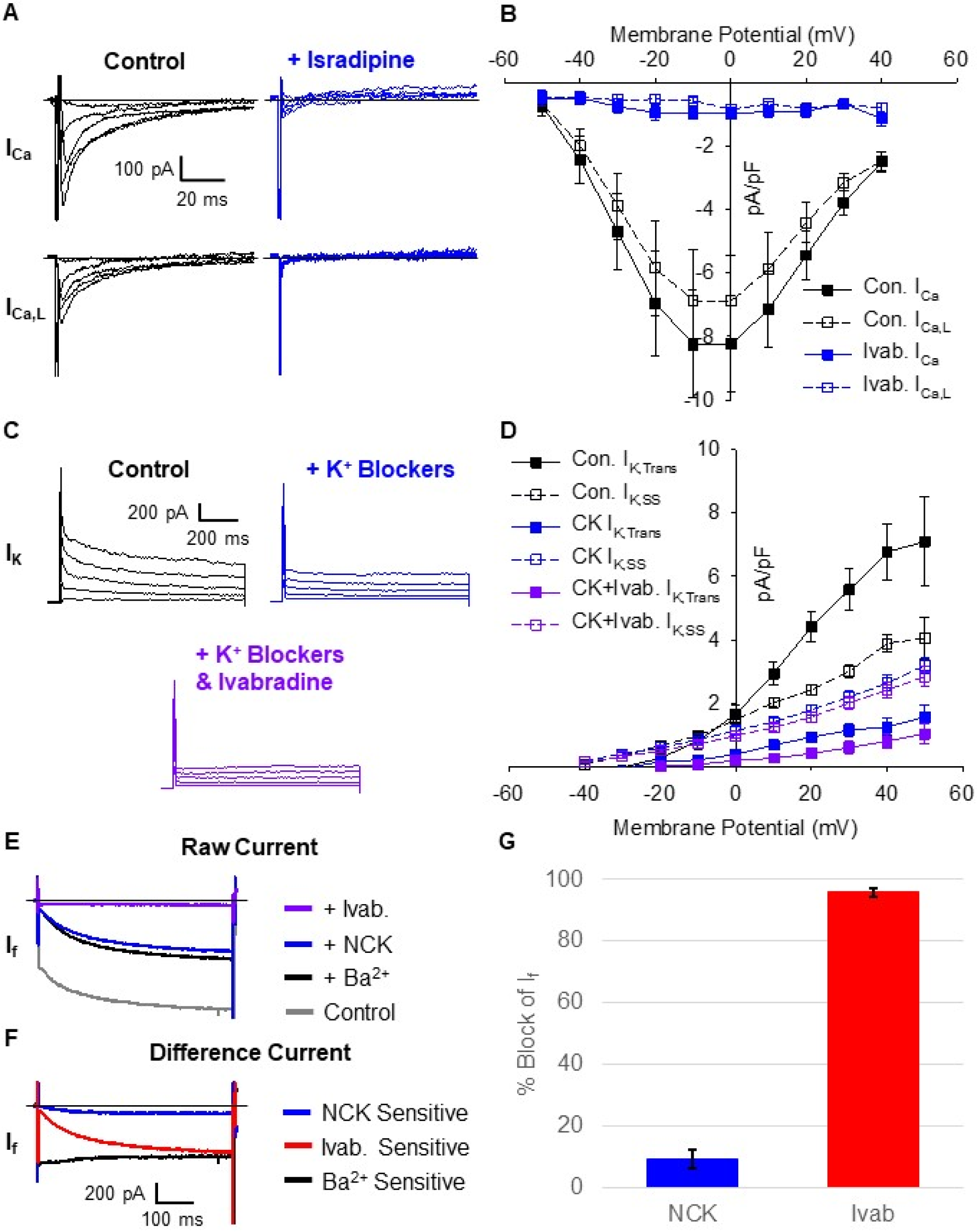
The NCK cocktail blocks ivabradine off-targets, but not I_f_. **A:** Representative total (*top*) and L-type (*bottom*) whole-cell Ca^2+^ currents recorded from sinoatrial node myocytes in control (*black*) and after perfusion of 3 µM isradipine (*blue*). **B:** Mean (± SEM) total (*closed*) and L-type (*open*) Ca^2+^ current density in control and after perfusion of 3 µM isradipine using the same colour scheme as **B**. **C:** Representative K^+^ currents in a SAM recorded in control conditions (*black*), after perfusion of a K^+^ channel blocker cocktail (1 mM BaCl_2_, 10 mM TEA, and 3 μM E-4031; *blue*), and after subsequent perfusion of the K^+^ channel blocker cocktail plus 30 µM ivabradine (*purple*). **D:** Average (± SEM) transient (*closed*) and steady-state (*open*) K^+^ current density in control conditions, after perfusion of a K^+^ channel blocker cocktail, and after subsequent perfusion of 30 µM ivabradine and a K^+^ channel blocker cocktail using the same colour scheme as **C**. **E:** Representative currents recorded from a murine SAM in response to hyperpolarizing voltage steps to -130 mV from a holding potential of -60 mV in Tyrode’s solution (*grey*), in Tyrode’s after application of 1 mM barium (*black*), the NCK blocker cocktail (*blue*), or the NCK cocktail with 30 µM ivabradine (*purple*). **F:** Representative difference currents sensitive to 1 mM barium compared to control (*black*), NCK cocktail compared to barium alone (*blue*), and the NCK cocktail with 30 µM ivabradine compared to the NCK cocktail alone (*red*). **G:** Average (± SEM) fraction of I_f_ blocked by the NCK cocktail (*blue*) or the NCK cocktail with 30 µM ivabradine (*red*) during pulses to -130 mV in murine SAMs. Horizontal lines indicate zero current. Number of replicates and details of statistical tests can be found in Table S2.

To determine whether the Ca^2+^ and K^+^ channel blockers had any effect on If, we next applied them together with the Na^+^ channel blocker tetrodotoxin (TTX; 30 μM) to isolated SAMs (Fig. 2E, F). 30 µM TTX was included in our pre-block cocktails because ivabradine has previously been shown to block Na^+^ currents (26) and 30 µM TTX inhibits both weakly TTX-sensitive Nav1.5 channels as well as highly TTX-sensitive Nav1.1 channels expressed in the murine sinoatrial node (42, 43). We found that the Na^+^-Ca^2+^-K^+^ (NCK) channel blocker cocktail blocked ∼10% of the inward current that was activated during 1 s hyperpolarizations to -120 mV (Fig. 2E,G; Table S2); however, the I_f_ amplitude after perfusion of the NCK cocktail did not differ significantly from the current after application of only the inward rectifier K^+^ channel blocker, barium (Tables S2), which has long been used to isolate I_f_ (21, 44). This indicates that TTX, isradipine, TEA, and E4031 exhibit no additional block of If. Subsequent addition of 30 μM ivabradine to the NCK cocktail significantly blocked I_f_ by ∼95% compared to barium alone (Fig. 2E,G; Table S2). Thus, the NCK blocker cocktail containing TTX, isradipine, TEA, barium, and E4031 can be used to effectively pre-block the off-target effects of ivabradine on Na^+^, Ca^2+^ and K^+^ currents in SAMs, with minimal effects on If. Unfortunately, the inhibition of total potassium current by the blocker cocktail was incomplete. The effect of ivabradine on the remaining potassium current was variable; in some cells ivabradine had no further effect on the remaining potassium current, while in other cells ivabradine inhibited a small component of current. Therefore, we analyzed I_f_ as defined by ivabradine-sensitive current only in the cells in which ivabradine had no effect on the steady-state potassium current remaining in the cocktail as measured by the reversal potential of I_f_ (see Methods).

### I_f_ flows both inward and outward during the cardiac cycle in SAMs

Having identified conditions for pharmacological isolation of I_f_ in SAMs, we next used the AP-clamp technique to record the native I_f_ flowing in response to APs recorded from the same cell. The experimental approach is illustrated in Fig. S2. In each cell, we first recorded spontaneous APs in current clamp mode without current injection in the presence of either 1 nM (control) or 1 µM (stimulated) of the β-adrenergic receptor agonist, isoproterenol. As expected, the AP firing rate was significantly faster in 1 µM isoproterenol compared to 1 nM isoproterenol (Table S3). From each of these recordings, a representative 10 s train of APs was integrated into a voltage protocol which appended square voltage steps to the end of the train of APs so that the degree of drug block could be monitored with the steps. Finally, repeated sweeps of the cell-specific AP-clamp voltage protocol were applied to each cell in voltage-clamp mode to elicit currents in a series of different solutions.

In each cell, AP-elicited currents were first recorded in the absence of blockers (but presence of 1 nM or 1 μM isoproterenol), then in response to wash-on of the NCK cocktail, and finally upon wash-on of the NCK cocktail plus 30 μM ivabradine (Fig. 3A, B). I_f_ was defined in these experiments as the ivabradine-sensitive difference current, which was calculated by subtracting the current after application of 30 μM ivabradine in the presence of the NCK cocktail from the current in the NCK cocktail alone (Fig. 3A, B). We confined the analysis of I_f_ during the firing cycle to cells in which ivabradine had no effect on the steady-state potassium current remaining in the blocker cocktail (see Methods). In those cells the reversal potential for the ivabradine-sensitive current did not differ from the reversal potential of steady-state I_f_ elicited by hyperpolarizing voltage steps in SAMs in 1 nM or 1 μM Iso (Fig. 3D; Table S3; P = 0.6973).

**Figure 3:**
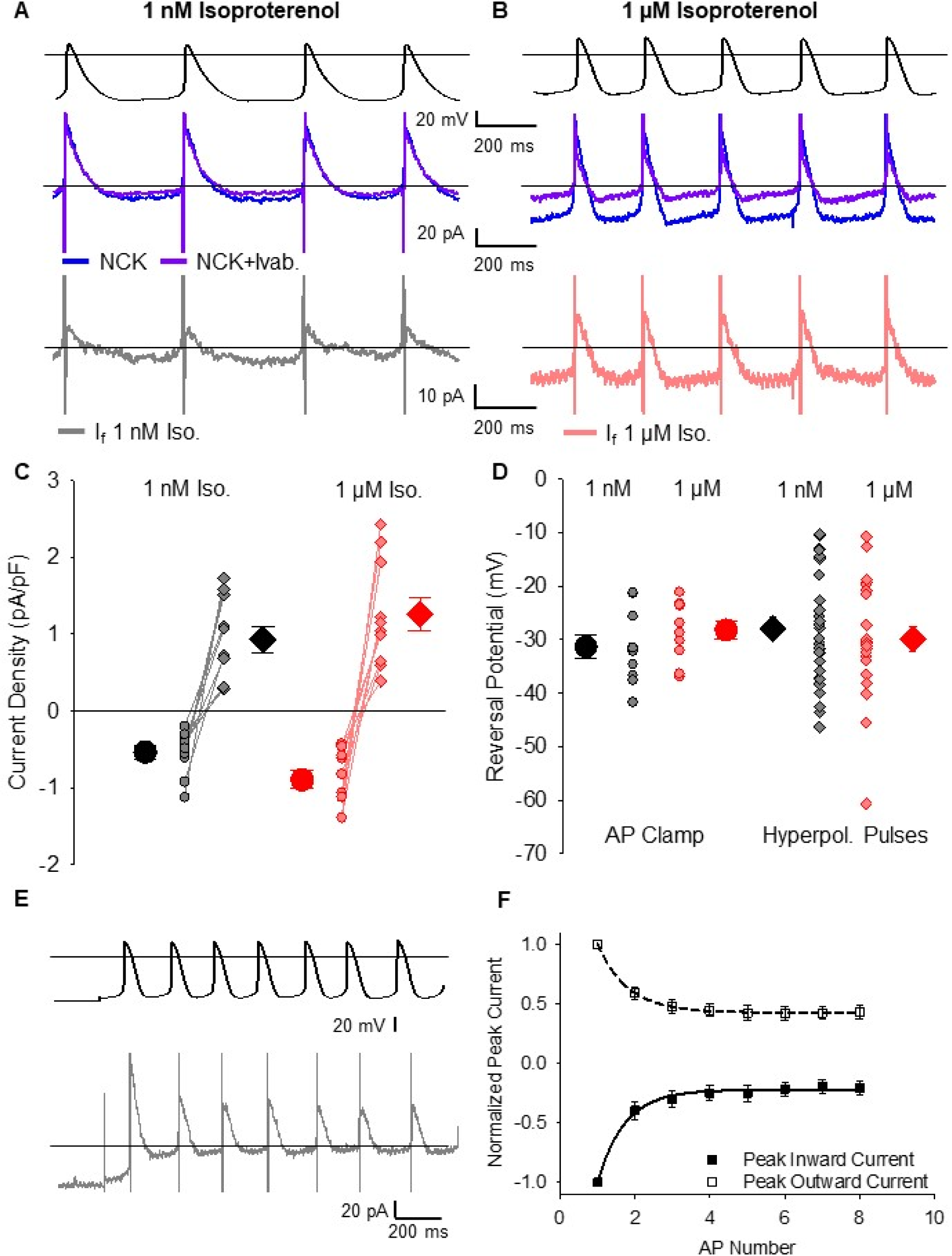
The funny current flows both inward and outward during sinoatrial Aps. **A-B:** Representative whole-cell currents (*middle*) elicited by APs recorded from the same cell (*top*) after the addition of the NCK cocktail (*blue*) or the NCK cocktail plus 30 μM ivabradine (*purple*). *Bottom*, the ivabradine-sensitive I_f_ difference current was determined as the current in NCK cocktail plus ivabradine minus the current in NCK cocktail alone in 1 nM isoproterenol (*grey*) or 1 μM isoproterenol (*pink*). **C:** Average (± SEM) peak inward (*circles*) and outward (*diamonds*) I_f_ current density in 1 nM isoproterenol (*black*) and 1 µM isoproterenol (*red*). Individual recordings in 1 nM isoproterenol (*grey*) or 1 µM isoproterenol (*pink*) are shown using the smaller symbols. **D:** Average (± SEM) reversal potential for ivabradine-sensitive current in 1 nM and 1 μM isoproterenol measured during AP waveforms (*circles*) and hyperpolarizing pulses (*diamonds*) using the same colour scheme as **C**. Number of replicates and details of statistical tests can be found in Table S3. **E:** Representative ivabradine-sensitive currents (*bottom*) in a sinoatrial myocyte in response to APs following a holding potential of -70 mV (*top*) in 1 nM isoproterenol. **F:** Average (± SEM) peak inward (*closed*) and peak outward (*open*) currents recorded in SAMs during the first 8 APs after holding at -70 mV in 1 nM isoproterenol (N=7). Average time courses were fit by a single exponential equation and are normalized to the peak current during the first AP in the same cell. The comparison of the time constants for inward and outward current decay was performed with a paired t-test (P = 0.7790). Horizontal lines indicate zero voltage or zero current levels.

I_f_ was evident as both an inward current at diastolic potentials and an outward current during the AP upstroke and repolarization phases. In the control condition of 1 nM isoproterenol, the inward component of the ivabradine-sensitive current peaked at an average of -18 pA and the outward current peaked at 29 pA (Table S3). As expected, 1 μM isoproterenol reduced the cycle length. 1 μM isoproterenol also significantly increased the peak inward I_f_ density during the AP, consistent with its expected enhancement of I_f_ (Fig. 3C; Table S3). Outward currents in 1 µM isoproterenol also tended to be larger than those in 1 nM isoproterenol (Fig. 3C), although the difference did not reach statistical significance.

We examined the correlation between the inward and outward ivabradine-sensitive currents by comparing their rates of change during a train of APs that was given following a holding potential of -70 mV (Fig. 3E). The relatively negative holding potential partially activated I_f_ before the train of APs, allowing us to measure the time-courses of decay in the peak outward and inward currents simultaneously as I_f_ deactivated to a new smaller steady state level during the train of APs. We found that the inward and outward ivabradine-sensitive currents decayed at similar rates (Fig. 3F; 200.4 ± 50.9 ms, n = 6 for inward; 191.4 ± 29.4 ms, n = 6 for outward; P = 0.7790), consistent with both originating from I_f_ channels with relatively slow relaxation kinetics as average membrane voltage is changed. Overall, these data establish that I_f_ is active during both the diastolic depolarization and repolarization phases of the sinoatrial AP.

### Current-voltage relationships of ivabradine-sensitive current during the cardiac cycle in SAMs

To further probe the attributes of the ivabradine-sensitive current, we next examined its current-voltage relationship. For each cell, the ivabradine-sensitive current was plotted as a function of the voltage of the AP command waveform. Current-voltage relationships were averaged for four consecutive APs (capacitive transients, which represent approximately 1 ms of data during the AP upstroke, were eliminated). We found that the current-voltage relationships for the ivabradine-sensitive current in both 1 nM and 1 μM isoproterenol were fairly linear at more negative potentials (Fig. 4A). The slope of the outward component of the current-voltage relationship appeared to be somewhat steeper than that of the inward component in many cells, reflecting the slight outward rectification of fully-activated I_f_ previously described (21, 45, 46). In some cells, there may also be a small amount of contamination of outward current by an ivabradine-sensitive outward current not fully blocked by the NCK cocktail.

**Figure 4:**
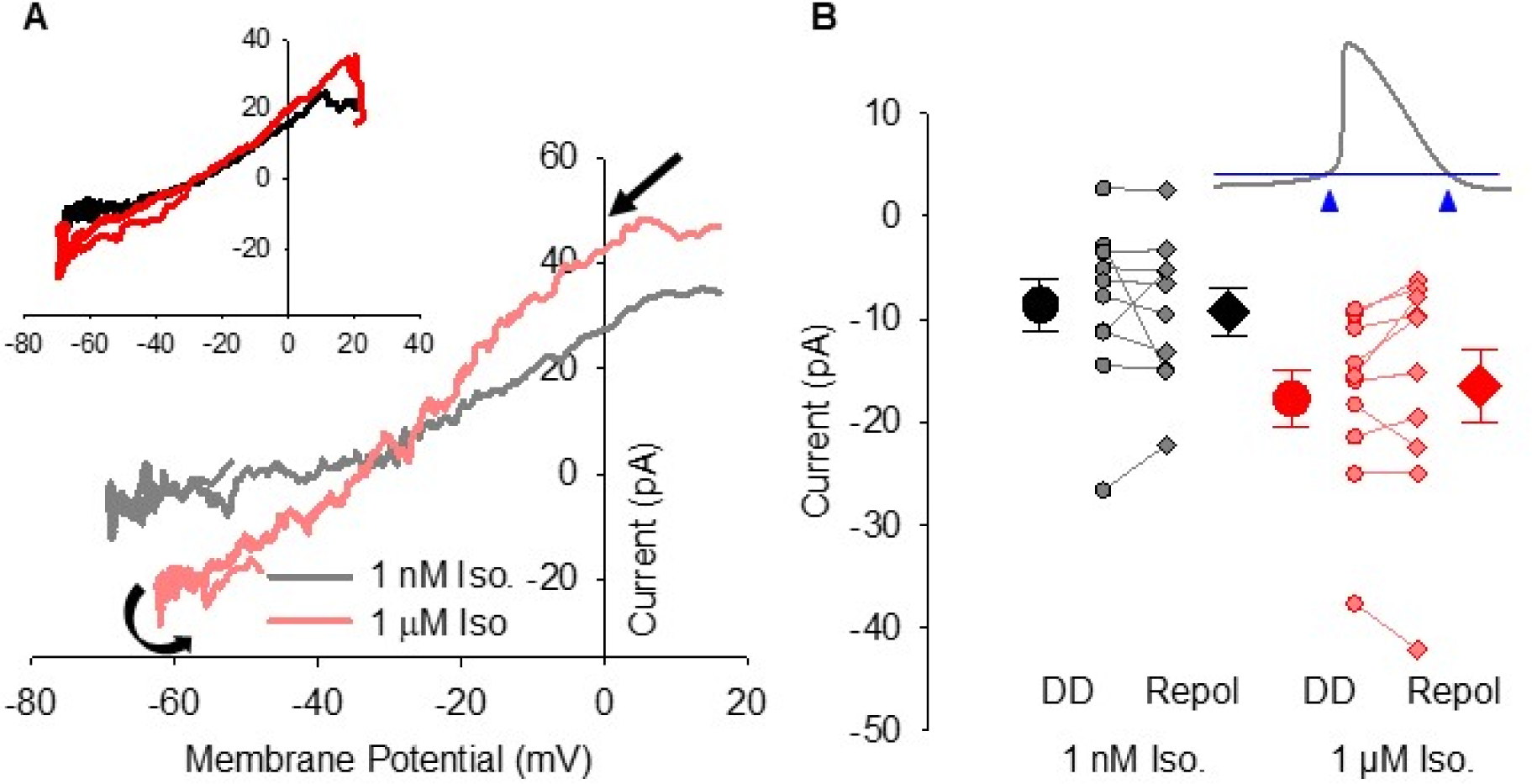
Current-voltage relationship and limited gating of I_f_ in SAMs. A: Representative current-voltage relationships of ivabradine-sensitive I_f_ from single cells in 1 nM isoproterenol (*grey*) or 1 µM isoproterenol (*pink*). Black arrows indicate the progression of the current-voltage relationship over time. **A *inset*:** Averaged current-voltage relationships of ivabradine-sensitive I_f_ in 1 nM isoproterenol (n = 10; *black*) or 1 µM isoproterenol (n = 10; *red*). **B:** Average (± SEM) I_f_ current amplitude recorded at -60 mV during the diastolic depolarization (DD; *circles*) and following AP repolarization (Repol; *diamonds*) in 1 nM isoproterenol (*black*) and 1 µM isoproterenol (*red*). Individual recordings in 1 nM isoproterenol (*grey*) or 1 µM isoproterenol (*pink*) are shown using the smaller symbols. **B *inset*:** Schematic showing points at which I_f_ was measured in panel **B**. Number of replicates and details of statistical tests can be found in Table S3.

The close tracking of the ivabradine-sensitive current to the AP waveform (Fig. 3A, B) and the fairly linear current-voltage relationships (Fig. 4A) suggest that I_f_ undergoes little voltage-dependent gating during the cardiac cycle in SAMs. To estimate the changes in I_f_ gating during the cardiac cycle, we compared the current amplitude at -60 mV during the latter part of the diastolic depolarization (just prior to the AP upstroke) to the current at -60 mV during the latter part of the AP repolarization (just following the AP downstroke) (Fig. 4B *inset*). We chose -60 mV as a test potential for these comparisons because it is sufficiently hyperpolarized relative to the reversal potential to allow accurate measurement of the small ivabradine-sensitive current and because it precedes the capacitive transient. We found that the average current amplitudes at -60 mV in SAMs did not differ between the depolarization and repolarization phases in either 1 nM or 1 µM isoproterenol (Fig. 4B; Table S3), consistent with the idea that there is little voltage-dependent gating of I_f_ during the AP in SAMs.

### I_f_ contributes substantially to the net current during both the diastolic depolarization and repolarization phases of the AP

We next estimated the contribution of I_f_ to the net inward and outward charge movement during the sinoatrial AP. The net current in each cell during the AP was calculated as the first derivative of membrane potential scaled by capacitance. For If, we extrapolated a linear fit to the I_f_ current-voltage relationship negative to -20 mV as a conservative estimate of the magnitude of outward I_f_ in the absence of any potential contamination (Fig. S3A, B). Comparison of I_f_ to the net current shows that I_f_ comprises a substantial fraction of the net charge moved during the sinoatrial AP. In 1 nM isoproterenol, I_f_ contributed 1.6 ± 0.35 pC (n = 10) of inward charge and 0.2 ± 0.05 pC (n = 10) of outward charge throughout each AP cycle (Fig. 5A; Table S3).

**Figure 5:**
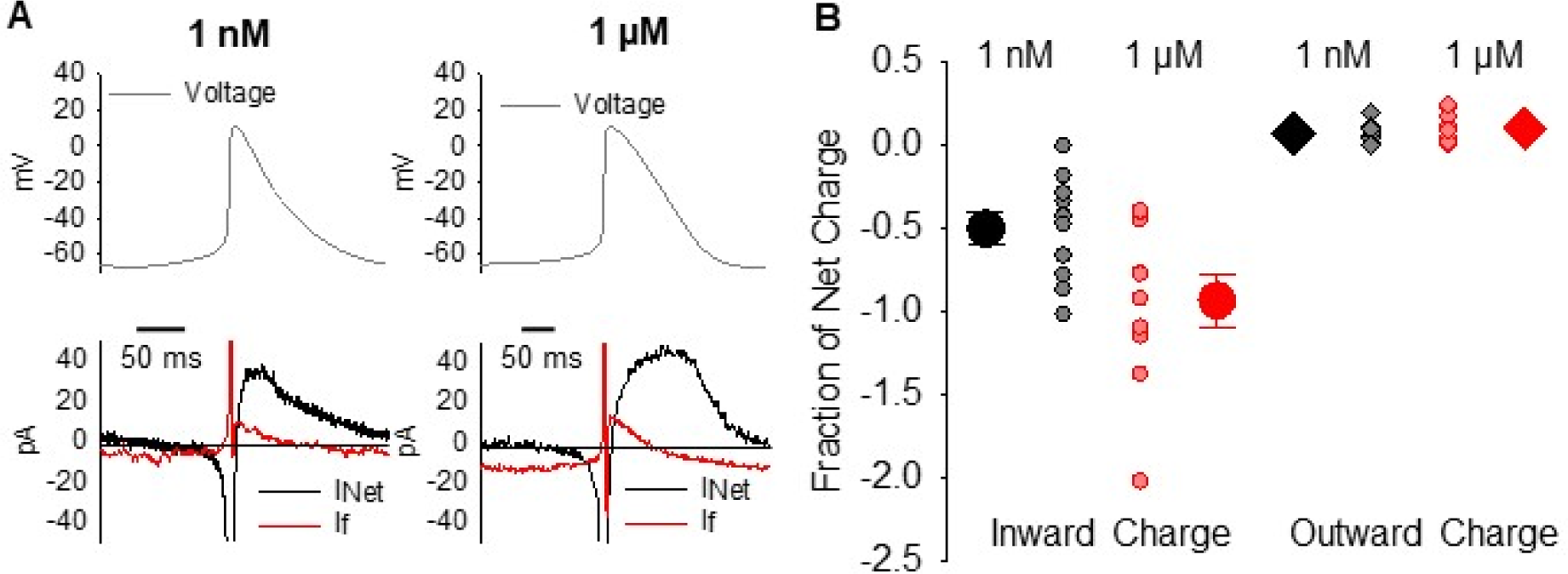
The funny current contributes substantial inward and outward charge movement during the sinoatrial node myocyte action potential. A: Representative calculated net current (*black*) and measured ivabradine-sensitive funny current (*red*) during a single sinoatrial AP (*top*) in 1 nM isoproterenol or in 1 µM isoproterenol. Horizontal lines indicate zero current. **B:** Average (± SEM) ivabradine-sensitive inward (*circles*) and outward (*diamonds*) charge movement as a fraction of the net charge during a single AP in 1 nM (*black*) and 1 μM (*red*). Individual recordings in 1 nM isoproterenol (*grey*) or 1 µM isoproterenol (*pink*) are shown using the smaller symbols. Number of replicates and details of statistical tests can be found in Table S3.

Importantly, this represents 50.2 ± 10.1% (n = 10) and 6.6 ± 1.9% (n = 10) of the net inward and outward charge movement, respectively (Fig. 5B). When compared to the ∼30 pF average cellular capacitance (Table S3), the ivabradine-sensitive I_f_ alone would be sufficient to depolarize the membrane by 45 mV and to repolarize the membrane by 6 mV during each AP in the absence of other conductances. Despite a shorter AP cycle length that decreases the amount of time I_f_ has to pass charge during each AP, 1 µM isoproterenol significantly increased the inward charge density moved by I_f_ by ∼40% (Table S3) and increased I_f_ as a fraction of the net inward current to 93.4 ± 16.0% (Fig. 5B; P = 0.0353). 1 μM Iso had no significant effect on outward ivabradine-sensitive charge movement, which remained about 7% of the net charge. In 1 µM isoproterenol, I_f_ contributes enough charge to depolarize the membrane by 85 mV and to repolarize it by 9 mV during each AP cycle. Ultimately, these data show that I_f_ contributes substantially to the charge movement during both the diastolic depolarization and repolarization phases of the sinoatrial AP. The significant increase in inward charge movement during diastole in response to 1 μM isoproterenol also indicates that I_f_ contributes to the βAR stimulated increase in AP firing rate.

### Limited voltage-dependent gating of HCN4 channels in HEK cells in response to sinoatrial APs

To better examine the properties of I_f_ during the sinoatrial AP, we next recorded currents passed by HCN4 channels expressed in HEK293 cells in response to APs recorded from SAMs (Fig. 6; Table S4). Control APs recorded in 1 nM Iso were used to evoke HCN4-mediated current in HEK cells without cAMP in the pipette, and APs recorded in 1 μM Iso were used in HCN4-expressing HEK cells where cAMP was present. Whole-cell currents were recorded before and after perfusion of 30 μM ivabradine in the presence of Ca^2+^ and K^+^ channel blockers (to block any endogenous currents in HEK cells) and in either the absence (Fig. 6A) or presence (Fig. 6B) of 1 mM cAMP in the recording pipette. The HCN4-generated current was determined as the ivabradine-sensitive difference current. In agreement with the above data from SAMs, HCN4 currents elicited by sinoatrial APs closely tracked the shape of the AP and had both inward and outward components (Fig. 6A, B). When plotted against the membrane potential, the HCN4 current was roughly linear throughout the duration of the AP with slight outward rectification (Fig. 6C). As expected, inclusion of 1 mM cAMP in the recording pipette significantly increased the inward and outward current densities during the AP and the slope of the current-voltage relationship (Fig. 6C; Table S4). Importantly, HCN4 currents in HEK cells exhibited little voltage-dependent gating during the AP. The IV relationships during the upstroke and repolarization phases of the AP were similar (Fig. 6C). Ivabradine-sensitive currents in HEK cells at -60 mV prior to the AP upstroke did not significantly differ from those measured at -60 mV during the AP downstroke in the absence of cAMP (Fig. 6D; Table S4). With 1 mM cAMP in the pipette, the current measured at -60 mV following repolarization was slightly (<5%) reduced in most cells compared to the current during the depolarization phase, consistent with a small amount of deactivation during the AP (Fig. 6D; Table S4). These results in HEK cells are in good agreement with our findings that native I_f_ passes substantial inward and outward current in response to the AP in SAMs and that there is little voltage-dependent gating of I_f_ during the sinoatrial AP.

**Figure 6:**
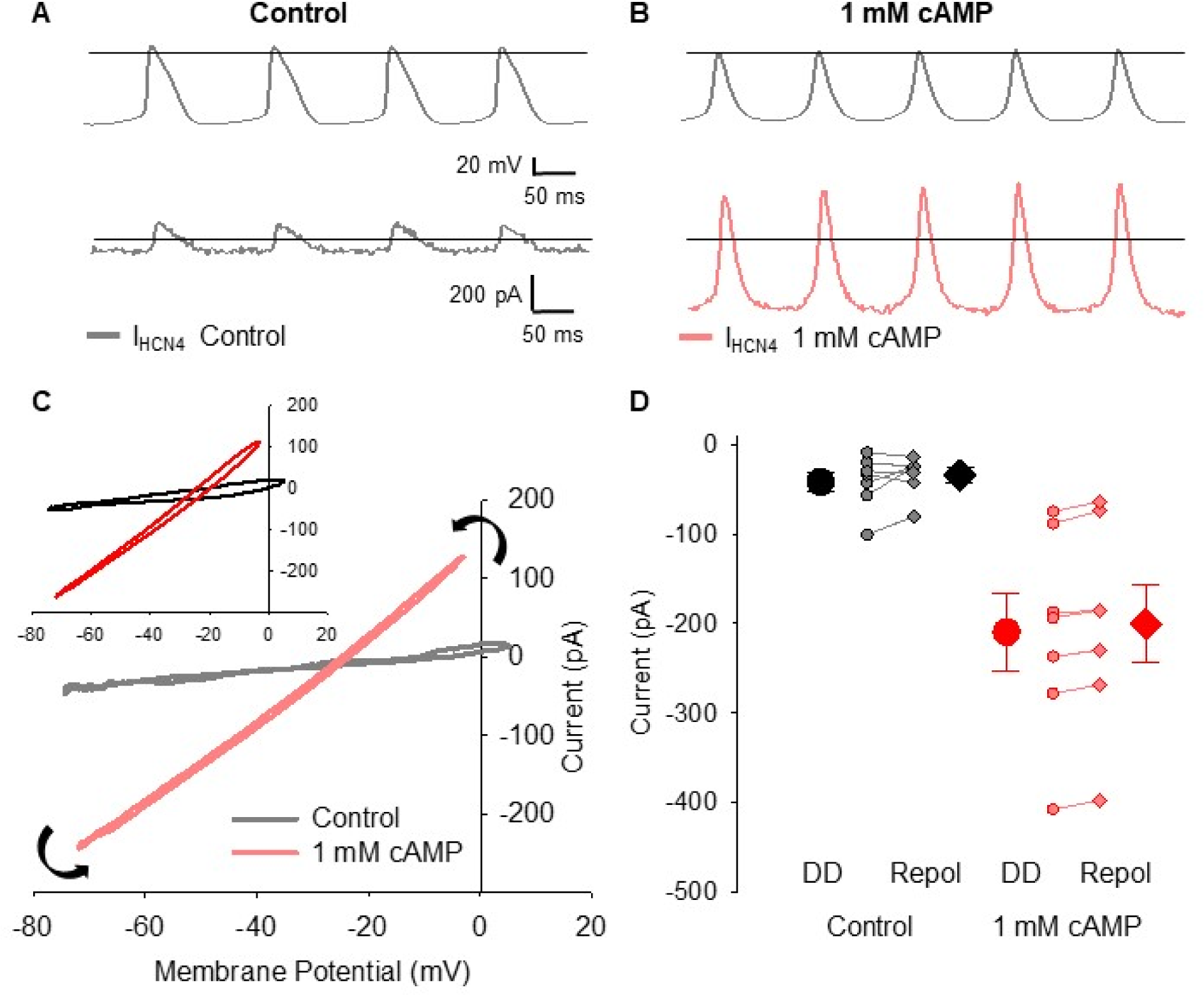
HCN4 channels in HEK cells pass a biphasic current with limited voltage-dependent gating in response to sinoatrial Aps. A: Representative ivabradine-sensitive HCN4 current in the absence of cAMP (*grey*) recorded from a HEK cell in response to APs recorded from a sinoatrial node myocyte in 1 nM isoproterenol (*top*). **B:** Representative ivabradine-sensitive HCN4 current with 1 mM cAMP in the intracellular solution (*pink*) recorded in a HEK cell in response to APs recorded from a sinoatrial node myocyte in 1 µM isoproterenol (*top*). Horizontal lines indicate zero voltage or zero current levels. **C:** Representative current-voltage relationships of ivabradine-sensitive HCN4 currents in control (*grey*) or 1 mM cAMP (*pink*). Black arrows indicate the progression of the current-voltage relationship over time. **C *inset*:** Average current-voltage relationships of ivabradine-sensitive HCN4 currents in control (*black*) and 1 mM cAMP (*red*). **D:** Average (± SEM) HCN4 current amplitude recorded at -60 mV during the latter diastolic depolarization (DD; *circles*) and AP repolarization (Repol; *diamonds*) in control (*black*) and 1 mM cAMP (*red*). Individual recordings in control (*grey*) or 1 mM cAMP (*pink*) are shown using smaller symbols. Number of replicates and details of statistical tests can be found in Table S4.

### Slow activation and deactivation of HCN4 underlies limited gating during the sinoatrial AP

We next used mathematical modelling to examine the biophysical properties of I_f_ that contribute to its persistent activation throughout the AP cycle in SAMs. We refined an existing model of I_f_ (47) to fit the currents elicited by square wave hyperpolarizing voltage protocols in murine SAMs in either 1 nM or 1 μM isoproterenol (Fig. S4). Sets of APs recorded from multiple cells in either 1 nM (n = 10) or 1 µM (n = 10) isoproterenol were then used to stimulate the models. For each simulation, the conductance of the model was scaled to match the I_f_ current measured in the cell from which the APs were recorded.

Although the simulated steady-state current-voltage relationships showed little current at potentials corresponding to the diastolic depolarization (Fig. S4C), both models produced currents with inward and outward components (Fig. 7A, B), similar to those recorded from SAMs or HCN4-expressing HEK cells (Figs. 3, 6). Simulated currents produced by the 1 µM model were larger than those produced by the 1 nM model (Fig. 7C inset, Fig. 7D; Table S5) and this difference in current amplitude persisted when the models were scaled to the same maximum conductance (Fig. S5A) or when the sets of APs used to stimulate the models were swapped (Fig. S5B), consistent with experimental observations that isoproterenol increases I_f_ amplitude primarily through changes to gating, not maximal conductance (21, 48, 49). The 1 nM model exhibited little or no voltage-dependent gating, as assessed by similar current-voltage relationships during the upstroke and downstroke (Fig. 7C) and similar current amplitudes at -60 mV before and after the AP (Fig. 7D; Table S5), consistent with the data acquired in SAMs and HCN4-expressing HEK cells (Figs. 4 & 6). However, the 1 μM isoproterenol model displayed a limited amount of gating by both measures, primarily in simulations with APs that had a more negative maximum diastolic potential (MDP) or larger AP amplitude (Fig. 7C & D).

**Figure 7:**
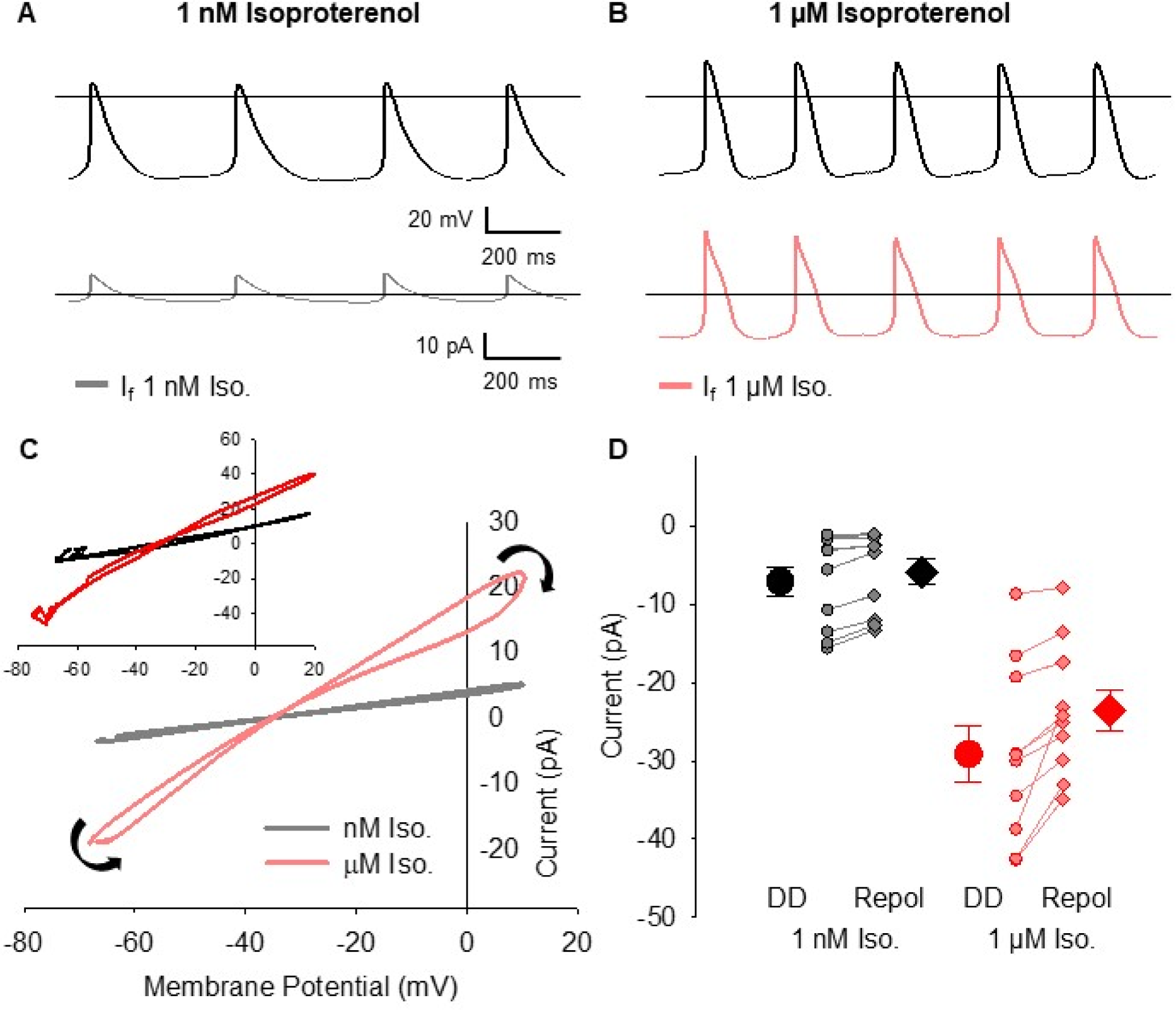
Simulations of the funny current produce a biphasic current with limited voltage-dependent gating in response to sinoatrial Aps. **A:** Simulated I_f_ generated by a model fit to data from 1 nM isoproterenol (*grey*) in response to APs recorded in 1 nM isoproterenol (*top*). **B:** Simulated I_f_ generated by a model fit to data from 1 μM isoproterenol (*pink*) in response to APs recorded in 1 µM isoproterenol (*top*). Horizontal lines indicate zero voltage or zero current levels. **C:** Representative current-voltage relationships of I_f_ simulated by the 1 nM model (*grey*) or the 1 μM model (*pink*) in response to sinoatrial APs recorded from single cells in the same condition. Black arrows indicate the progression of the current-voltage relationship over time. **C *inset*:** Average current-voltage relationships of I_f_ simulated by the 1 nM model (*black*) or the 1 μM model (*red*) in response to the full sets of sinoatrial APs recorded in 1 nM or 1 μM isoproterenol. **D:** Average (± SEM) simulated I_f_ current amplitude at -60 mV during the latter diastolic depolarization (DD; *circles*) and AP repolarization (Repol; *diamonds*) in response to recorded sinoatrial APs simulated using the 1 nM isoproterenol model (*black*) or the 1 µM isoproterenol model (*red*). Individual simulations in response to different 1 nM (*grey*) or 1 μM (*pink*) APs are shown by the smaller symbols. Number of replicates can be found in Table S5.

We hypothesized that the lack of substantial gating of I_f_ during the AP is caused by the relatively slow rates of activation and deactivation of the sinoatrial HCN4 isoform. To test this hypothesis, we compared the response of the 1 nM model of I_f_ to those of previously published models of HCN1 (50) and HCN2 (51), which activate and deactivate up to an order of magnitude more rapidly than HCN4 (6, 52–56). Compared to HCN4, which activates with a time constant of ∼450 ms at -140 mV, HCN1 and HCN2 activation time constants at -140 mV are ∼30 ms and ∼184 ms, respectively (6). Similarly, at -50 mV the deactivation time constants in HCN1 and HCN2 (∼125 ms and ∼600 ms, respectively) are far more rapid than that of HCN4 (∼1.6 s) (17).

In contrast to the 1 nM I_f_ model, both the HCN1 and HCN2 models showed prominent activation during the AP cycle, as seen by figure-eight shaped current-voltage relationships (Fig. 8A). Indeed, the HCN1 model, which has the fastest kinetics and most depolarized voltage-dependence, completely deactivated during the AP upstroke and contributed little outward current during the AP repolarization phase. HCN2, which has activation and deactivation rates more similar to those of HCN4, showed considerable outward current during the upstroke, but was mostly deactivated during the AP downstroke.

**Figure 8:**
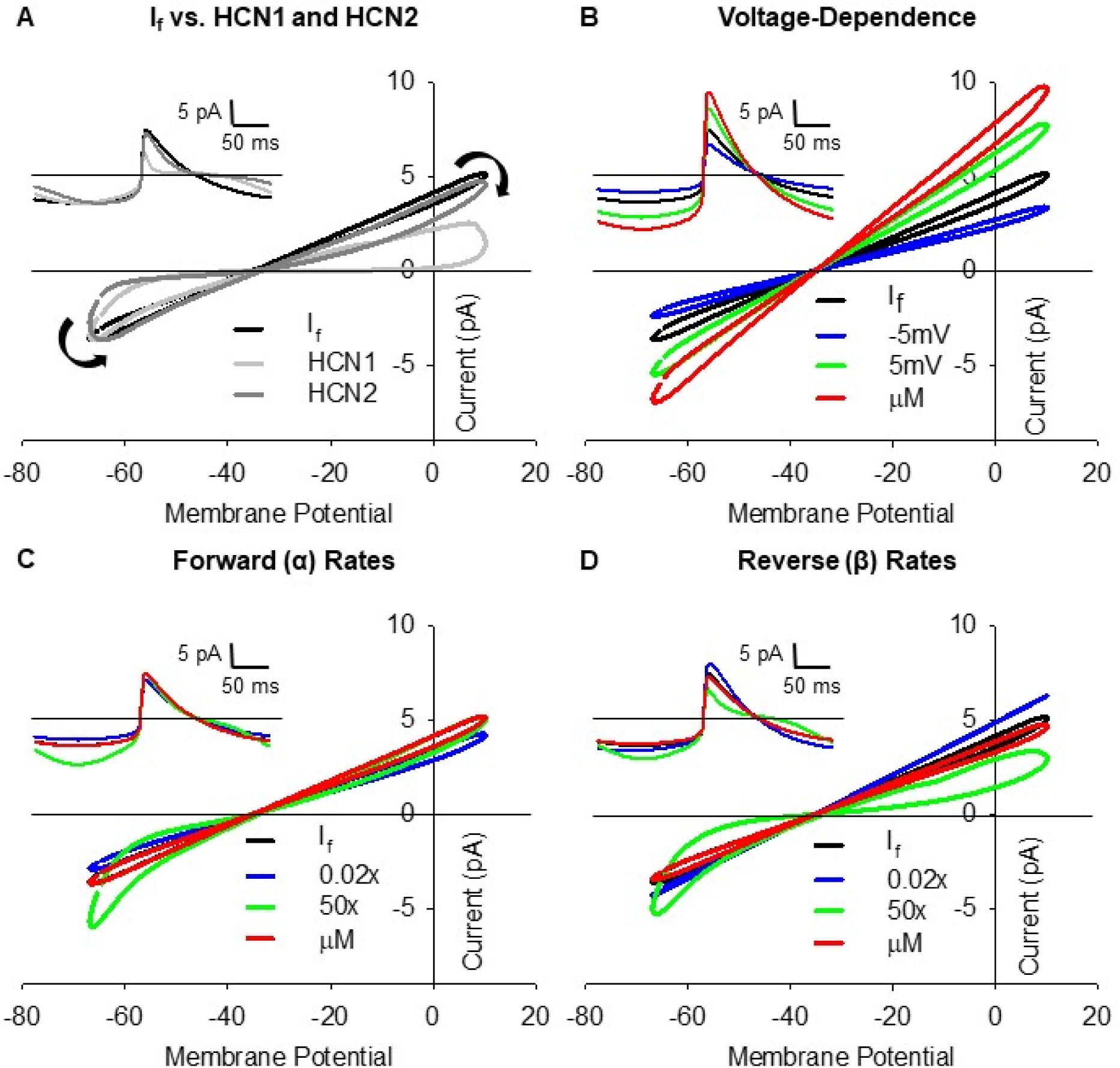
Linear funny current is due to slow activation and deactivation rates. **A:** Simulated current-voltage relationships in response to a representative sinoatrial AP recorded in 1 nM isoproterenol using the base 1 nM isoproterenol model of I_f_ (*black*), a model of HCN1 (*light grey*) (50), or a model of HCN2 gating (*dark grey*) (51). Black arrows indicate the progression of the current-voltage relationship over time. **A *inset***: Simulated currents during single sinoatrial APs using the base 1 nM isoproterenol model of If, the HCN1 model or the HCN2 model. **B-D:** Current-voltage relationships of simulated I_f_ in response to pre-recorded sinoatrial APs using the base 1 nM isoproterenol model (*black*), the 1 µM isoproterenol model (*red*), or the 1 nM model with altered activation voltage-dependence (**B**), forward (α) rates of activation (**C**), or reverse (β) rates of activation (**D**). Activation voltage-dependence (V_1/2_) was shifted by -5 (*blue*) or +5 mV (*green*). The α and β rates of the activation gate were slowed by 50x (*blue*) or sped up 50x (*green*). **B-D *insets***: Simulated I_f_ during single sinoatrial APs using the base 1 nM isoproterenol model, the 1 µM isoproterenol model, or models with altered activation voltage-dependence (**B *inset***), α rates (**C *inset***), or β rates (**D *inset***). Colour schemes in **B-D *insets*** are the same as the main panels. Horizontal lines indicate zero current.

To elucidate how the individual gating parameters impact the current-voltage relationships of If, we compared the base 1 nM model to the 1 nM model in which the three individual components of the model were changed independently (voltage-dependent activation, the forward rate (α) of the activation transition, and the reverse rate (β) of the activation transition (Fig. 8B-D)). We found that 5 mV depolarizing or hyperpolarizing shifts in the voltage-dependence of activation or a 15 mV depolarizing shift corresponding to stimulation by 1 μM isoproterenol primarily increased or decreased the slope of the IV relationship, without much change in the amount of deactivation during the AP (∼20% in all cases; Fig. 8B). In contrast, a model in which the forward rate was sped up by 50 times led to prominent I_f_ activation during the diastolic depolarization (Fig. 8C). And a model with the reverse rate sped up 50 times showed both I_f_ activation during the diastolic depolarization and I_f_ deactivation during the AP upstroke and repolarization phases (Fig. 8D). Changing the α and β rates of the 1 nM model to those in the 1 µM model or slowing them by 50x did not substantially alter channel gating during the AP (Fig. 8C and 8D). Thus, the persistent activation of I_f_ during the sinoatrial AP appears to be mainly determined by the slow activation and deactivation rates of the underlying HCN4 channels.

## Discussion

Despite decades of research dedicated to defining the ionic currents that underlie cardiac pacemaking in SAMs, little consensus has emerged on how I_f_ contributes to the generation of spontaneous APs (9, 12, 57, 58). We here establish a new conceptual framework for the role of I_f_ as a driver of oscillatory electrical activity in SAMs by showing that it flows both inward and outward during the AP. Our results specifically reveal that: 1) I_f_ is active throughout the sinoatrial AP and passes both inward and outward current; 2) I_f_ contributes a substantial portion of the net depolarizing and repolarizing charge movement during the sinoatrial AP; 3) there is limited voltage-dependent gating of I_f_ during the cardiac cycle in SAMs; and 4) the slow rates of activation and deactivation of the sinoatrial HCN4 channel isoform contribute to the persistent activation of I_f_ throughout the cardiac cycle in SAMs.

### I_f_ in SAMs drives membrane potential oscillations in both directions during the AP

Most studies of I_f_ have focused on its role as an inward current during the diastolic depolarization phase of the sinoatrial AP (9, 12, 14–16). While mathematical models and some previous studies have suggested that I_f_ may also conduct outward current in SAMs (18–20, 59, 60), our data are the first experimental demonstration that I_f_ is active throughout the entire AP cycle (Fig. 3-5). This behavior was confirmed in our AP-clamp recordings of HCN4-mediated currents in HEK cells (Fig. 6) and our AP clamp simulation of a mathematical model of I_f_ parameterized using traditional square-wave protocols (Fig. 7). Together the data indicate that I_f_ serves a critical role in the generation of spontaneous activity in the cardiac pacemaker by driving membrane potential oscillations in both directions. It follows that I_f_ must contribute to shaping both the diastolic depolarization and repolarization phases of the sinoatrial AP. This unique function of I_f_ arises as a result of both its reversal potential and its slow kinetics; the reversal potential of ∼ -30 mV confers driving force at both extremes of the voltage excursion of the sinoatrial AP (roughly -60 to +10 mV) whilst the slow rates of activation and deactivation conspire to keep the hyperpolarization-activated channels open throughout the entire AP cycle.

### Limited voltage-dependent gating due to slow kinetics of HCN4

Strikingly, our data show that there is little voltage-dependent activation or deactivation of I_f_ during the cardiac cycle in SAMs, either at resting AP firing rates (in 1 nM isoproterenol) or at the higher firing rates seen in response to βAR stimulation (in 1 μM isoproterenol). The lack of appreciable gating was recapitulated by heterologously-expressed HCN4 channels and by mathematical models of I_f_ (Figs. 6 & 7). Our subsequent modelling (Fig. 8) suggests that the voltage-independent nature of I_f_ recorded in AP clamp experiments is mainly determined by the intrinsically slow rates of activation and deactivation of HCN4 compared to other HCN isoforms. Our findings help resolve the paradox of how such a slowly-activating current can contribute to the diastolic depolarization of sinoatrial cells (12, 13) by showing that the equally slow deactivation rate results in a conductance that is activated throughout the entirety of the AP cycle.

An important consequence of this persistent activation is that I_f_ contributes a substantial fraction of the total charge movement in SAMs that belies its relatively small peak current amplitude and low open probability. Indeed, the peak amplitude of I_f_ during the cardiac cycle in SAMs is only ∼10-30 pA, consistent with its negative range of voltage activation relative to the voltages of the sinoatrial AP (e.g., Fig. S4C). These currents correspond to activation of only a small fraction of the total HCN channels in the cell. Based on an average maximal conductance of 11.79 nS measured during hyperpolarizing voltage steps (Fig. S4), a maximum of ∼2% of HCN channels are open during the diastolic depolarization in 1 nM isoproterenol and ∼5% in 1µM isoproterenol. Yet, despite the small peak current amplitudes and low open probability, I_f_ contributes 50% of the net inward charge (≈3.1 nC) and 6.5% of the net outward charge (≈3.1 nC) during the AP (Fig. 5B) due to its persistent activity. These results provide a mechanistic framework for numerous qualitative observations from mutation, blocker and simulation studies that illustrate that I_f_ is critical for pacemaker function.

### Increased I_f_ during diastole contributes to the fight-or-flight response

The lack of gating during the cardiac cycle does not mean that voltage-dependent gating of I_f_ is irrelevant for the physiological function of HCN4 channels. Our data also show that the shift in voltage-dependence and changes in kinetics in response to βAR stimulation yield a new gating equilibrium with larger currents and increased charge movement throughout the AP cycle (Figs. 3-7). During βAR stimulation, inward charge movement through I_f_ increases nearly 40%, driven by increased channel open probability in response to cAMP and PKA phosphorylation (Fig. 8B) (3, 61, 62).

To date, knockout and knockdown of HCN4 in mice have consistently shown that I_f_ is necessary to stabilize basal AP firing rates in SAMs (9, 63–66). However, the data have been less clear on the role of I_f_ during β-adrenergic stimulation. SAMs isolated from inducible knockouts of HCN4 in adult mice respond normally to isoproterenol in some lines (65, 66), but have a decreased response in others (63). And the inducible deletion of cyclic nucleotide binding via the HCN4-573X mutant reduces exercise-induced maximal heart rate and decreases maximal AP firing rate of isolated cells in response to isoproterenol, but does not fully eliminate β-adrenergic stimulated increases in SAM firing and heart rate (64). Importantly, interpretation of all these models is complicated by the presence of residual If, possibly due to expression of other HCN channel isoforms (65). Furthermore, this lack of clarity regarding the contribution of I_f_ may reflect altered regulation of other mechanisms of heart rate regulation when HCN4 is knocked-out. While our data do not imply that I_f_ is the only driver of fight-or-flight increases in heart rate, they demonstrate conclusively that β-adrenergic stimulation increases the I_f_ amplitude and its relative contribution to the inward charge movement during the diastolic depolarization, consistent with a role for I_f_ in the fight-or-flight increase in heart rate as has long been suggested (3).

### Limitations of blockers

Some of the historic difficulty in evaluating the role of I_f_ in pacemaking stems from the imperfection of available blockers. For example, Cs^+^ has long been used to block If, however, Cs^+^ also blocks inward rectifier K^+^ channels and alters gating of Kv11.1 (hERG) (28–30). Moreover, the Cs^+^-sensitive current in SAMs underestimates I_f_ because Cs^+^ block of inward I_f_ is relieved at more depolarized membrane potentials and because Cs^+^ does not block outward current through HCN channels (46). Substantial off-target effects have similarly been reported for other HCN channel blockers, including ZD7288 (27, 33), zatebradine (31, 32, 36, 37), and cilobradine (32, 67). We used ivabradine in the present study as it has relatively low arrhythmogenic potential at the therapeutic concentration of 3 µM (8, 36, 37, 68) and it is widely used as a research tool to study HCN channel currents in native tissues (38, 64, 69–71). However, our experiments required fairly complete block of If, and we found that at the concentration required to achieve about 90% block of I_f_ in SAMS (30 μM), ivabradine exhibited substantial off-target block of voltage-gated Ca^2+^ and K^+^ currents (Fig. 1). This is consistent with previous reports (26, 34, 35) and with its increased arrhythmogenic potential at higher concentrations (39). Despite the off-target effects, we were able to measure I_f_ as the ivabradine-sensitive current in SAMs by pre-blocking Na^+^, K^+^, and Ca^2+^ currents (Fig. 2), similar to methods used to isolate AP-elicited currents mediated by Kv3 channels in DRG neurons (72) and the sequential blocker application approach used to isolate currents in ventricular myocytes (73).

### Self-AP Clamp Compensates for Cellular Heterogeneity

Measurement of I_f_ during the sinoatrial AP is further complicated by the large variability between AP waveforms in sinoatrial myocytes (21–23, 74). Thus, instead of applying “representative” APs to all cells, we customized the approach for each cell, by applying APs recorded from the same cell as the voltage command, as has been previously reported in DRG neurons (72) and ventricular myocytes (75). This approach has the advantage that the AP waveform is precisely matched to the ion channel expression in each cell. Our data highlight the AP-clamp technique as a powerful tool to record non-equilibrium properties of ion channels in native tissues and assess their contributions to cellular excitability. In the present study, we used the AP-clamp technique to define I_f_ in SAMs. In future work, a similar approach can be used to isolate other currents in SAMs and, ultimately, to refine and validate mathematical models of pacemaking.

### Conclusions

The data presented herein demonstrate that I_f_ has both inward and outward components during the cardiac cycle in mouse SAMs. The slow gating kinetics of the HCN4 isoform and a reversal potential of -30 mV allow I_f_ to stay active throughout the sinoatrial AP and contribute not only to the diastolic depolarization but also to the repolarization phase of the AP. Although the current amplitude is quite small, I_f_ comprises a large fraction of the net charge movement in both inward and outward directions. Our findings extend the long-standing notion of I_f_ as an inward current during the diastolic depolarization by showing that I_f_ contributes to oscillatory electrical activity in SAMs by driving membrane potential back towards its reversal potential in both directions.

## Materials and Methods

### Animal Ethics and Sinoatrial Node Dissociation

This study was carried out in accordance with the US Animal Welfare Act and the National Research Council’s *Guide for the Care and Use of Laboratory Animals* and was conducted according to a protocol that was approved by the University of Colorado-Anschutz Medical Campus Institutional Animal Care and Use Committee.

Six-to eight-week old male C57BL/6J mice were obtained from Jackson Laboratories (Cat. #000664). Animals were anesthetized by isofluorane inhalation and euthanized under anesthesia by cervical dislocation. Hearts were excised and bathed in warmed (35°C) modified Tyrode’s solution (in mM: 140 NaCl, 5.4 KCl, 1.2 KH_2_PO_4_, 5 HEPES, 5.55 glucose, 1 MgCl_2_, 1.8 CaCl_2_; pH adjusted to 7.4 with NaOH) containing 10 U/mL heparin. The ventricles were removed and the sinoatrial node was dissected from the atrial tissue, as defined by the borders of the crista terminalis, the interatrial septum, and the superior and inferior vena cavae, as described in previous studies (24, 76).

The sinoatrial node was cut into three to five strips, which were digested at 35°C for in a low Ca^2+^ modified Tyrode’s solution (in mM: 140 NaCl, 5.4 KCl, 1.2 KH_2_PO_4_, 5 HEPES, 18.5 glucose, 0.066 CaCl_2_, 50 taurine, 1 mg/ml BSA; pH adjusted to 6.9 with NaOH) containing either 4.75 U elastase (Worthington Biochemical) and 3.75 mg Liberase TM (Roche) for 10-12 minutes or 1064 U type II collagenase (Worthington), 9 U elastase (Worthington), and 652 µg type XIV protease (Sigma) for 30 min. Following enzymatic digestion, SAMs were dissociated by mechanical trituration with a fire-polished glass pipette (∼2 mm diameter) for 5 - 8 min at 35°C in a modified KB solution (in mM: 100 K^+^-glutamate, 10 K^+^-aspartate, 25 KCl, 10 KH_2_PO_4_, 2 MgSO_4_, 20 taurine, 5 creatine, 0.5 EGTA, 20 glucose, 5 HEPES, and 0.1% BSA; pH adjusted to 7.2 with KOH). Ca^2+^ and BSA were gradually introduced to the cell suspension in 5 steps over the course of 22 min to a final concentration of 1.8 mM Ca. Cells were stored in KB solution at room temperature for up to 10 hours before electrophysiological recordings.

### Electrophysiology

Data were acquired at 5-20 kHz and low-pass filtered at 1 kHz using an Axopatch 1D or 200B amplifier, Digidata 1322a or 1440a A/D converter, and ClampEx software (Molecular Devices). Bath and perfusing solutions were maintained at 35°C for all recordings with a feedback-controlled temperature controller (TC-344B, Warner Instruments). Reported data have been corrected for liquid junction potentials between the pipette and extracellular solutions for all voltage-clamp recordings. Fast (pipette) capacitance was compensated in all voltage-clamp recordings. Recording pipettes were pulled from borosilicate glass (VWR International) using a Sutter Instruments P-97 horizontal puller. Pipette resistances ranged from 1.5 to 5 MΩ.

### Sinoatrial Myocyte Current Clamp Recordings

An aliquot (∼50 ul) of the isolated SAM suspension was transferred to a glass bottomed recording chamber and cells were allowed to settle for ∼5 minutes before perfusion with extracellular solution. Recording Tyrode’s solution (in mM: 140 NaCl, 5.4 KCl, 5 HEPES, 5 Glucose, 1 MgCl_2_, 1.8 CaCl_2_; pH adjusted to 7.4 with NaOH) containing 1 nM isoproterenol was used to record AP firing rates. Pipettes were filled with an internal solution composed of (in mM): 110 K-aspartate, 20 KCl, 1 MgCl_2_, 5 EGTA, 5 Mg-ATP, 5 creatine phosphate, and 5 HEPES; pH adjusted to 7.2 with KOH. After a gigaohm seal and whole-cell configuration were established, cells were lifted off the bottom of the recording chamber and the amplifier was set to current clamp mode with zero injected current. 30 s of stable AP firing was recorded followed by up to 2.5 min of AP firing with perfusion of 30 µM ivabradine in Tyrode’s solution. The average cell firing rate over the last 10 s of every 30 s period was calculated using ParamAP software (77).

### Sinoatrial Myocyte Square-Wave Voltage-Clamp Recordings

Whole-cell Ca^2+^ currents were recorded in a K^+^ current-blocking extracellular solution (in mM: 130 tetraethylammonium chloride, 2 CaCl_2_, 1 MgCl_2_, 10 4-aminopyridine, and 10 HEPES; pH adjusted to 7.4 with CsOH) with a pipette solution consisting of 130 mM CsCl, 1 mM MgCl_2_, 10 mM Hepes, 10 mM EGTA, 4 mM Mg-ATP, and 0.1 mM NaGTP, with the pH adjusted to 7.2 with CsOH as described previously (24). Total Ca^2+^ current was measured in response to depolarizing steps between -50 to +40 mV from a holding potential of -90 mV. L-type Ca^2+^ currents were measured in response to depolarizing steps between -50 to +40 mV from a holding potential of -60 mV where T-type current is inactivated. T-type currents were estimated by subtracting the L-type Ca^2+^ current from total Ca^2+^ current in each cell. Leak subtraction was performed using a P/4 protocol. Ivabradine and isradipine block of currents were assessed following perfusion of 30 μM ivabradine or 3 μM isradipine as indicated.

Whole-cell K^+^ currents were measured in response to depolarizing steps between -40 to +50 mV from a holding potential of -50 mV in recording Tyrode’s extracellular solution containing 1 nM isoproterenol with an intracellular solution consisting of 135 mM KCl, 1 mM MgCl_2_, 4 mM Mg-ATP, 0.1 mM Na-GTP, 5 mM EGTA, 5 mM HEPES, and 6.6 mM Na-phosphocreatine, with the pH adjusted to 7.2 with KOH. Cells were continuously perfused at 35 ℃ with Tyrode’s solution containing 3 μM isradipine to block voltage-gated Ca^2+^ currents. Steady-state current was measured at the end of the depolarizing step, while the transient current was measured by subtracting steady-state current from the peak outward current during a depolarizing step. Inhibition by ivabradine and the K-blocker cocktail were assessed following perfusion of 30 μM ivabradine and/or 10 mM TEA, 1 mM barium, and 1 µM E4031 as indicated.

Whole-cell I_f_ recordings for use in modelling were measured in recording Tyrode’s solution containing either 1 nM or 1 µM isoproterenol as indicated. Pipettes were filled with K-aspartate intracellular solution. I_f_ was measured as the time-dependent current elicited by 3 s steps to potentials between -60 and -150 mV from a holding potential of -50 mV. Activation voltage-dependence was determined by calculating conductance from time-dependent inward currents as G=I/V-V_rev_, where G is conductance, I is current, V is the step potential, and V_rev_ is the I_f_ reversal potential (-30 mV; (61). The normalized conductance for each cell was fit with a single Boltzmann equation to calculate the midpoint and slope of the relationship. Time constants and the relative amplitudes of the fast and slow components of I_f_ activation and deactivation were measured by double-exponential fits. Deactivation was measured during 3 s steps to potentials between -50 and +50 mV following a 1 s step to -150 mV.

### Sinoatrial Myocyte AP-Clamp Recordings

For AP-clamp recordings, SAMs were patch-clamped in the whole-cell configuration and constantly perfused at 35°C with recording Tyrode’s extracellular solution containing 1 nM or 1 µM isoproterenol as indicated. Pipettes were filled with an internal solution composed of (in mM): 140 K-aspartate, 13.5 mM NaCl, 1.8 MgCl_2_, 0.1 EGTA, 4 Mg-ATP, 0.3 GTP (tris salt), 14 creatine phosphate, and 9 HEPES, pH 7.2 with KOH. Spontaneous APs were first recorded in the whole-cell current clamp configuration and then 10 s of the AP recording were included in the command voltage protocol in voltage-clamp experiments to isolate the specific currents flowing during the AP (Fig. S2). The protocol was first used to record ionic currents in the absence of compensation for cellular capacitance to ensure that the amplifier currents were near zero (75).

To isolate If, we subtracted currents immediately before and after applying 30 μM ivabradine, while in the constant presence of 1 mM BaCl_2_, 10 mM TEA, 1 μM E-4031, 30 μM TTX and 3 μM isradipine. I_f_ block was monitored during hyperpolarizing steps to -90 mV or -120 mV. To account for incomplete block of I_f_ by ivabradine, the subtracted currents during AP-clamp were scaled for the measured block achieved in the hyperpolarizing pulses from the same cell. In some cells, the current inhibited by ivabradine included a component of potassium current that remained in the blocker cocktail (Fig. S3C), as evident by current evoked by a step to 0 mV after a prepulse to -40 mV (which deactivates If) or by reversal potentials of the ivabradine-sensitive current that were substantially negative to the reversal potential for I_f_ under the ionic conditions (∼31 mV, Fig. 3D, Fig. S3D). Analysis was restricted to cells in which no contamination by potassium current was evident.

### HEK Cell Electrophysiology

HEK 293 cells stably expressing HCN4 were grown at 37°C at 5% CO_2_ in high glucose DMEM with L-glutamine, supplemented with 10% FBS, 100 U/mL penicillin, and 100 µg/ml streptomycin (all from Thermo Fisher Scientific, Waltham, MA). To maintain the HEK-HCN4 stable line, the media was further supplemented with 200 μg/mL Hygromycin B (Invivogen, San Diego, CA). HEK 293 cells were negative for mycoplasma infection. Testing for mycoplasma infection was performed at the Molecular Biology Core Facility in the Barbara Center for Childhood Diabetes at the University of Colorado Anschutz Medical Campus.

24 hours prior to experiments cells were seeded on protamine-coated glass coverslips. Recordings were performed in recording Tyrode’s extracellular solution with 1-2 MΩ borosilicate glass pipettes filled with K-Aspartate intracellular solution containing 0 or 1 mM cAMP as indicated. All recordings were performed 35°C. After a gigaohm seal and whole-cell configuration were established, we recorded HCN4 currents using previously recorded murine sinoatrial APs as a voltage command. For 0 mM and 1 mM cAMP recordings, APs recorded at 1 nM isoproterenol or 1 µM isoproterenol were used, respectively. HCN4 currents were recorded by subtracting currents immediately before and after applying 30 μM ivabradine, while in the presence of a “CK” blocker cocktail containing: 1 mM BaCl_2_, 10 mM TEA, 3 μM E-4031, and 3 μM isradipine. HCN4 current amplitude and current-voltage relationships were calculated from an average of 3 consecutive APs from each cell.

### Modelling

We modified an existing Hodgkin-Huxley type model of I_f_ in murine SAMs (47) and manually tuned the parameters to fit our experimental datasets obtained in SAMs in the presence of 1 nM or 1 μM isoproterenol. In both conditions, the total I_f_ current is calculated as the sum of the inward sodium (I_f,Na_) and the outward potassium (I_f,K_) components scaled by the relative ion permeabilities through the channel pore (Equation 1):

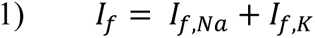

The two individual components were calculated as follows (Equations 2-4):

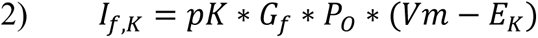

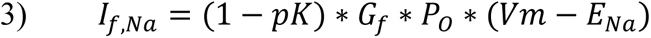

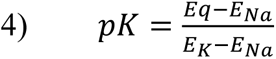

Where Vm is the membrane potential, Gf is the maximal I_f_ conductance, Po is the channel open probability, pK is the permeability to potassium relative to sodium, E_q_ is the I_f_ equilibrium potential, and E_Na_ and E_K_ are the Nernst potentials of Na^+^ and K^+^, respectively. In our novel formulation, the activation/deactivation gate n determining PO is controlled by both fast and slow processes nf and ns (Equations 5-6).

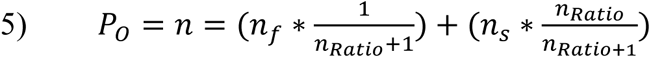

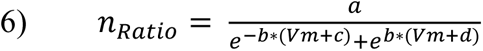

Where n_Ratio_ indicates the distribution of channels between slow and fast gating modes ns and nf at a given membrane potential, and was obtained fitting the ratio between the amplitudes of the slow and fast components estimated from biexponential fitting of current traces (Fig. S2D). Changes over time in nf and ns are calculated as follows (Equations 7-11):

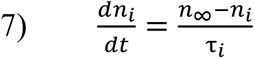

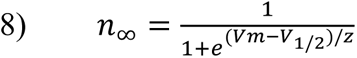

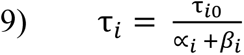

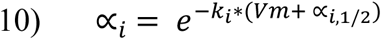

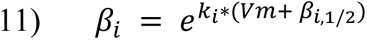

Where n_∞_ is the steady-state activation common to both nf and ns (Fig. S2C), and τ_i_ is the time constant of activation or deactivation at a given voltage for either the fast (Fig. S2E) or slow (Fig. S2F) component of the activation gate. α_i_ and β_i_ are the activation and deactivation rates for either the fast or slow component of the activation gate. Values of parameters used in Equations 1-11 are reported in Table 1.

To model I_f_ during the sinoatrial AP, sets of AP waveform recordings from the 1 nM and 1 μM isoproterenol AP-clamp experiments were used to stimulate the 1 nM and 1 μM isoproterenol models, respectively. Current-voltage relationships were calculated from the average of 3 consecutive APs. Model conductances were scaled for individual runs such that the current amplitude at either -90 mV or -120 mV was the same as that experimentally recorded in the SAM from which the voltage protocol was derived.

The model of HCN2 gating uses the equations from the non-cAMP bound state of the model proposed by Wang *et al.* (51). The HCN1 model uses the gating scheme proposed by Battefeld *et al.* (50). The maximal conductances of the HCN1 and HCN2 models were scaled such that the maximal inward currents during AP clamp simulations were similar to the 1 nM I_f_ model to facilitate comparisons of altered gating.

All models were simulated in the Spyder Python environment using a forward Euler method with a 200 µs time step.

### Analysis and Statistics

All analysis was performed in Clampfit 10.7 (Molecular Devices), with SigmaPlot 12.0 (SYSTAT Software) used for plotting. All statistical analysis was performed in JMP 14 (SAS Institute). AP firing rate in response to 30 μM ivabradine was evaluated with a repeated measures ANOVA with time following application of ivabradine as the independent variable. Comparisons to firing rate in the absence of ivabradine were performed with a Tukey post-hoc test. Blockade of Ca^2+^ currents by ivabradine or isradipine were evaluated using paired t-tests with the respective drug as the independent variable. Ivabradine and/or K^+^-blocker cocktail inhibition of transient and steady-state K^+^ currents were evaluated with a repeated measures ANOVA with the presence of specific blockers as the independent variable followed by a Tukey post-hoc test for individual comparisons. To compare currents or charge movement of experimentally measured I_f_ and HCN4 in response to isoproterenol or cAMP, respectively, we used student’s t-tests. Paired t-tests were used to compare I_f_ or HCN4 current amplitude at -60 mV during the AP upstroke to the amplitude at -60 mV during AP repolarization in the same cells. Significance was determined at P < 0.05. All means, standard errors of the mean, and N-values are provided in in text or in supplementary tables S1-S3.

## Abbreviations

AP: Action potential
βAR: β adrenergic receptor
HCN: Hyperpolarization-activated cyclic nucleotide-sensitive channel
I_f_: Funny current
SAM: Sinoatrial node myocyte

## Acknowledgements and Funding Sources

This work was funded by grants from the National Institutes of Health (HL 088427 to CP; NS036855 to BPB; R01HL131517, R01HL141214, P01HL141084, and Stimulating Peripheral Activity to Relieve Conditions Grant OT2OD026580 (to EG); R00HL138160 (to SM), and by a postdoctoral fellowship from the American Heart Association (19POST34380777 to CHP). The authors would like to acknowledge Dr. Christian Rickert for his insight and guidance on experimentation and the manuscript and Dr. Emily Sharpe for her work designing the AP clamp protocol and blocker series.

## Supplementary Material

**Supplementary Table 1:** Off-target effects of ivabradine in SAMs.

| | Control Peak Density<br>(pA/pF) | 3 μM Ivabradine Peak<br>Density (pA/pF) | 30 μM Ivabradine Peak Density<br>(pA/pF) |
| --- | --- | --- | --- |
| **I<sub>f</sub>** | -22.86 ± 3.84 (7) | -13.23 ± 3.33 (7)<br>P = 0.0157 vs. Control | -1.01 ± 0.37 (7)<br>P < 0.0001 vs. Control |
| **I<sub>Ca</sub>** | -11.36 ± 1.29 (5) | N/A | -7.50 ± 0.61 (5)<br>P = 0.0236 vs. Control |
| **I<sub>Ca,L</sub>** | -10.01 ± 1.06 (5) | N/A | -5.70 ± 0.81 (5)<br>P = 0.0046 vs. Control |
| **I<sub>Ca,T</sub>** | -1.35 ± 0.29 (5) | N/A | -1.80 ± 0.74 (5)<br>P = 0.6329 vs. Control |
| **I<sub>K,Trans</sub>** | -9.57 ± 1.50 (5) | N/A | -5.21 ± 1.65 (5)<br>P = 0.0391 vs. Control |
| **I<sub>K,SS</sub>** | 2.78 ± 0.22 (5) | N/A | 1.89 ± 0.19 (5)<br>P = 0.0191 vs. Control |

I<sub>f</sub>: Funny Current; I<sub>Ca</sub>: Total Ca<sup>2+</sup> current; I<sub>Ca,L</sub>: L-type Ca<sup>2+</sup> current; I<sub>Ca,T</sub>: T-type Ca<sup>2+</sup> current;
I<sub>K,Trans</sub>: Transient K<sup>+</sup> current; I<sub>K,SS</sub>: Steady-state K<sup>+</sup> current.
Peak Ca<sup>2+</sup>, K<sup>+</sup>, and I<sub>f</sub> currents were measured at 0 mV, +40 mV, and -130 mV, respectively. Data are averages ± SEM with number of replicates in parentheses. Tests for statistical differences in peak I<sub>f</sub> density were performed using repeated measures ANOVAs of currents recorded with either 3 μM or 30 μM ivabradine matched to currents recorded in control conditions (1 mM BaCl<sub>2</sub>) in the same cell. Tests for statistical differences in peak current density of Ca<sup>2+</sup> and K<sup>+</sup> currents were performed using paired t-tests of currents recorded with 30 μM ivabradine matched to currents recorded in control conditions in the same cell.

**Supplementary Table 2:** Ca^2+^ and K^+^ blocker effects in SAMs.

| | Control Peak Density<br>(pA/pF) | Ca<sup>2+</sup> and/or K<sup>+</sup> Blockers<br>Peak Density (pA/pF) | Ca<sup>2+</sup> and/or K<sup>+</sup> Blockers with 30 μM<br>Ivabradine Peak Density (pA/pF) |
| --- | --- | --- | --- |
| **I<sub>Ca</sub>** | -8.22 ± 1.70 (9) | -0.96 ± 0.16 (9)<br>P = 0.0022 vs. Control | N/A |
| **I<sub>Ca,L</sub>** | -6.86 ± 1.59 (9) | -0.57 ± 0.09 (9)<br>P = 0.0030 vs. Control | N/A |
| **I<sub>Ca,T</sub>** | -5.08 ± 0.91 (9) | -0.46 ± 0.06 (9)<br>P = 0.00057 vs. Control | N/A |
| **I<sub>K,Trans</sub>** | 6.78 ± 0.87 (9) | 1.25 ± 0.27 (9)<br>P < 0.0001 vs. Control | 0.81 ± 0.24 (9)<br>P = 0.7173 vs. Blockers |
| **I<sub>K,ss</sub>** | 3.90 ± 0.26 (9) | 2.63 ± 0.24 (9)<br>P < 0.0001 vs. Control | 2.41 ± 0.26 (9)<br>P = 0.3127 vs. Blockers |
| **I<sub>f</sub>** | -17.25 ± 4.60 (8) | -15.76 ± 4.32 (8)<br>P = 0.8726 vs. Control | -0.73 ± 0.34 (8)<br>P = 0.0002 vs. Control |

I<sub>Ca</sub>: Total Ca<sup>2+</sup> current; I<sub>Ca,L</sub>: L-type Ca<sup>2+</sup> current; I<sub>Ca,T</sub>: T-type Ca<sup>2+</sup> current; I<sub>K,Trans</sub>: Transient K<sup>+</sup> current; I<sub>K,ss</sub>:
Steady-state K<sup>+</sup> current; I<sub>f</sub>: Funny Current.

Peak Ca^2+^, K^+^, and I_f_ currents were measured at 0 mV, +40 mV, and -130 mV, respectively. Data are averages ± SEM with number of replicates in parentheses. Tests for statistical differences in peak current density of Ca^2+^ currents were performed using paired t-tests of currents recorded with isradipine matched to currents recorded in control conditions in the same cell. Tests for statistical differences in peak current density of K^+^ currents were performed using repeated measures ANOVAs of currents recorded with a K^+^ blocker cocktail in the absence or presence of ivabradine matched to currents recorded in control conditions in the same cell. Tests for statistical differences in peak I_f_ density were performed using repeated measures ANOVAs of currents recorded with the NCK blocker cocktail in the absence or presence of ivabradine matched to currents recorded in control conditions (1 mM BaCl_2_) in the same cell.

**Supplementary Table 3:** Ivabradine-sensitive current during AP clamp in SAMs.

| | 1 nM Isoproterenol | 1 μM Isoproterenol | 1 μM vs. 1 nM |
| --- | --- | --- | --- |
| **Firing Rate (bpm)** | 268 ± 212 (10) | 340 ± 26 (10) | P = 0.0474 |
| **Cellular Capacitance (pF)** | 33.7 ± 2.6 (10) | 30.1 ± 2.54 (10) | P = 0.3336 |
| **Inward Current** |  |  |  |
| *Peak Current - (pA)* | -17.7 ± 2.9 (10) | -26.7 ± 4.4 (10) | P = 0.1046 |
| *Peak Current Density - (pA/pF)* | -0.54 ± 0.09 (10) | -0.90 ± 0.12 (10) | P = 0.0240 |
| *Charge movement (pC)* | -1.61 ± 0.35 (10) | -2.64 ± 0.64 (10) | P = 0.1713 |
| *Charge Density - (pC/pF)* | -0.045 ± 0.009 (10) | -0.085 ± 0.014 (10) | P = 0.0283 |
| *Conductance - (nS)* | 0.25 ± 0.05 (10) | 0.57 ± 0.09 (10) | P = 0.0058 |
| **Outward Current** |  |  |  |
| *Peak Current - (pA)* | 29.1 ± 5.2 (10) | 36.9 ± 6.5 (10) | P = 0.3597 |
| *Peak Current Density - (pA/pF)* | 0.93 ± 0.17 (10) | 1.26 ± 0.22 (10) | P = 0.2584 |
| *Charge movement (pC)* | 0.35 ± 0.08 (10) | 0.56 ± 0.17 (10) | P = 0.2692 |
| *Charge Density - (pC/pF)* | 0.011 ± 0.003 (10) | 0.018 ± 0.005 (10) | P = 0.2658 |
| *Conductance - (nS)* | 0.49 ± 0.07 (10) | 0.56 ± 0.11 (10) | P = 0.6027 |
| **Corrected Outward Current** |  |  |  |
| *Peak Current - (pA)* | 10.5 ± 2.6 (10) | 17.9 ± 5.0 (10) | P = 0.2064 |
| *Peak Current Density - (pA/pF)* | 0.34 ± 0.09 (10) | 0.62 ± 0.18 (10) | P = 0.1895 |
| *Charge movement (pC)* | 0.19 ± 0.05 (10) | 0.29 ± 0.11 (10) | P = 0.3890 |
| *Charge Density - (pC/pF)* | 0.006 ± 0.002 (10) | 0.009 ± 0.003 (10) | P = 0.3497 |
| *Conductance - (nS)* | 0.15 ± 0.05 (10) | 0.27 ± 0.07 (10) | P = 0.2200 |
| **Reversal Potential (mV)** | -31.3 ± 2.2 (10) | -28.2 ± 1.8 (10) | P = 0.2769 |
| **-60 mV Current** |  |  |  |
| *Upstroke - (pA)* | -8.6 ± 2.5 (10) | -17.8 ± 2.7 (10) | |
| *Downstroke - (pA)* | -9.3 ± 2.3 (10) | -16.5 ± 3.6 (10) | |
|  | P = 0.6826 | P = 0.2982 |  |

All data are averages ± SEM with number of replicates in parentheses. Corrected outward currents are calculated from extrapolated linear fits to data from potentials more negative than -20 mV to eliminate contamination by unblocked K^+^ currents in some cells. Tests for statistical differences between 1 nM and 1 µM isoproterenol were performed using unpaired t-tests. Tests for statistical differences between current at -60 mV between the upstroke and downstroke of the action potential were performed using paired t-tests.

**Supplementary Table 4:** Ivabradine-sensitive current in response to AP Clamp in HCN4-expressing HEK cells.

|  | 0 mM cAMP | 1 mM cAMP | 0 mM vs. 1 mM |
| --- | --- | --- | --- |
| **Inward Current** |  |  |  |
| Peak Current - (pA) | -62.7 ± 14.3 (7) | -264.1 ± 54.9 (7) | P = 0.0040 |
| Peak Current Density - (pA/pF) | -2.05 ± 0.71 (7) | -5.23 ± 1.07 (7) | P = 0.0136 |
| Charge movement (pC) | -5.51 ± 1.33 (7) | -23.54 ± 4.91 (7) | P = 0.0041 |
| Charge Density - (pC/pF) | -0.195 ± 0.078 (7) | -0.466 ± 0.095 (7) | P = 0.0474 |
| Conductance - (nS) | 1.75 ± 0.42 (7) | 7.07 ± 1.45 (7) | P = 0.0046 |
| **Outward Current** |  |  |  |
| Peak Current - (pA) | 26.6 ± 9.6 (7) | 135.6 ± 38.1 (7) | P = 0.0169 |
| Density - (pA/pF) | 0.71 ± 0.16 (7) | 2.58 ± 0.65 (7) | P = 0.0160 |
| Charge movement (pC) | 0.31 ± 0.13 (7) | 1.02 ± 0.29 (7) | P = 0.0489 |
| Charge Density - (pC/pF) | 0.007 ± 0.002 (7) | 0.019 ± 0.005 (7) | P = 0.0482 |
| Conductance - (nS) | 0.75 ± 0.23 (7) | 5.11 ± 1.05 (6) | P = 0.0011 |
| **Reversal Potential (mV)** | -21.2 ± 4.8 (7) | -22.9 ± 3.0 (7) | P = 0.7590 |
| **-60 mV Current** |  |  |  |
| Upstroke - (pA) | -42.0 ± 11.4 (7) | -209.7 ± 43.2 (7) |  |
| Downstroke - (pA) | -34.6 ± 8.3 (7) | -200.8 ± 43.5 (7) |  |
|  | P = 0.2594 | P = 0.0011 |  |

**Supplementary Table 5:** Simulated funny current in response to AP waveforms.

| | 1 nM Isoproterenol | 1 μM Isoproterenol |
| --- | --- | --- |
| **Inward Current** |  |  |
| *Peak - (pA)* | $-7.5 \pm 2.7$ (10) | $-15.9 \pm 2.3$ (10) |
| *Charge - (pC)* | $-1.23 \pm 0.38$ (10) | $-4.36 \pm 0.80$ (10) |
| **Outward Current** |  |  |
| *Peak - (pA)* | $18.9 \pm 5.9$ (10) | $33.6 \pm 4.2$ (10) |
| *Charge - (pC)* | $0.26 \pm 0.07$ (10) | $0.85 \pm 0.16$ (10) |
| **-60 mV Current** |  |  |
| *Upstroke - (pA)* | $-7.1 \pm 1.9$ (10) | $-29.1 \pm 3.6$ (10) |
| *Downstroke - (pA)* | $-5.9 \pm 1.6$ (10) | $-23.6 \pm 2.7$ (10) |

All data are averages ± SEM with number of replicates in parentheses.

**Figure S1:**
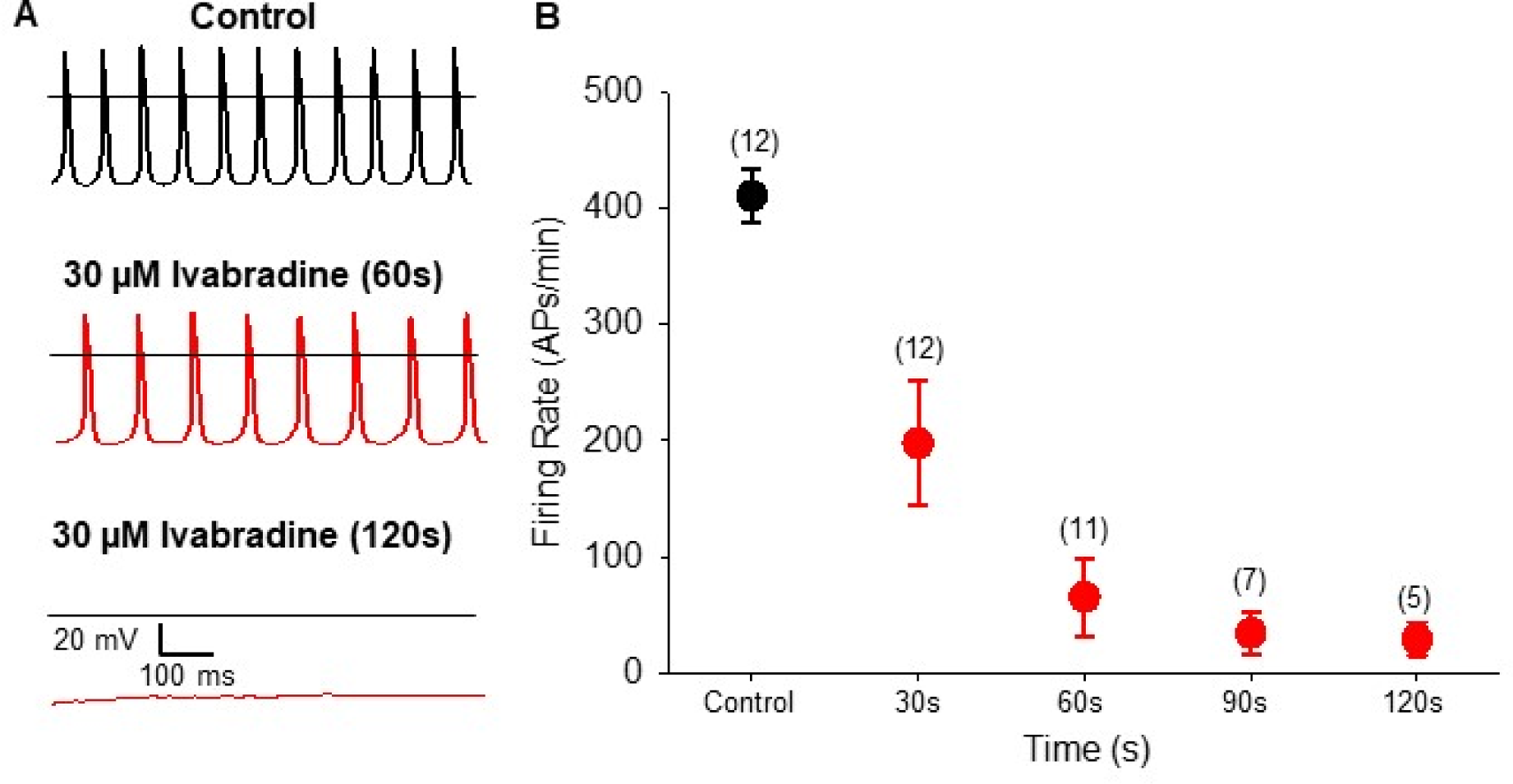
Ivabradine slows AP firing in SAMs. **A:** Representative APs recorded from a sinoatrial myocyte in control Tyrode’s solution containing 1 nM isoproterenol (*top*) or following 60 s (*middle*) or 120 s (*bottom*) of perfusion with 30 µM ivabradine. Horizontal lines indicate 0 mV. **B:** Average AP firing rates (± SEM) before (*black*) or at different time points during 30 µM ivabradine perfusion (*red*). Number of observations for each time point are given in parentheses.

**Figure S2:**
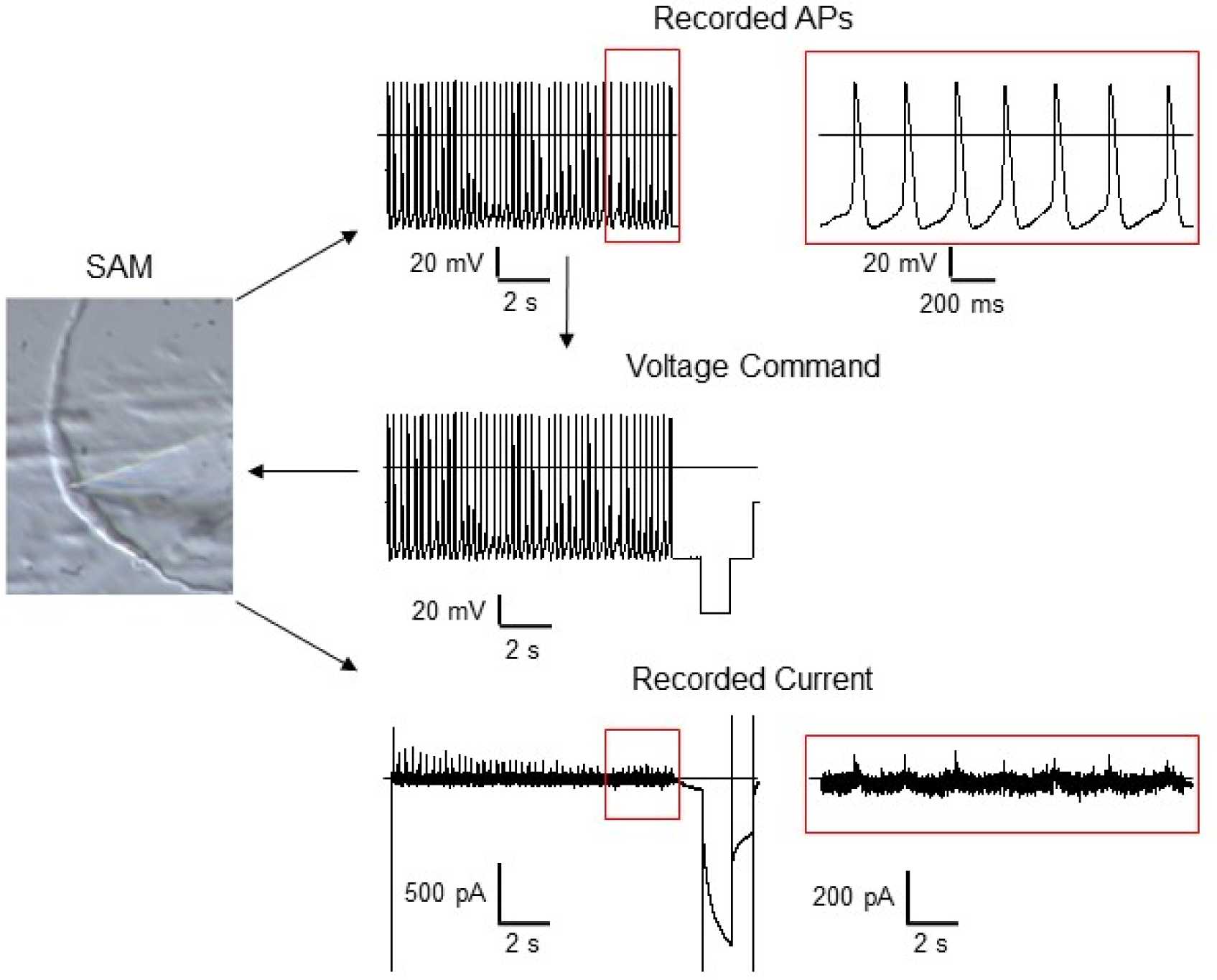
AP clamp methodology. Schematic of the workflow used in AP clamp experiments. Spontaneous APs were first recorded in current clamp mode without injected current (I = 0) from isolated SAMs in either 1 nM or 1µM isoproterenol (*top* and *top inset*). 10 s of APs were then integrated into a voltage protocol that also included square voltage pulses at the end of each sweep so that drug block could be monitored over time (*middle*). Repeated sweeps of this voltage protocol were then applied to the same myocyte in voltage-clamp mode. As expected, the current elicited by APs from the same cell is near zero in the control Tyrode’s solution without any drugs (and without whole-cell capacitance compensation) (*bottom* and *bottom inset*). Horizontal lines indicate zero voltage or zero current levels.

**Figure S3:**
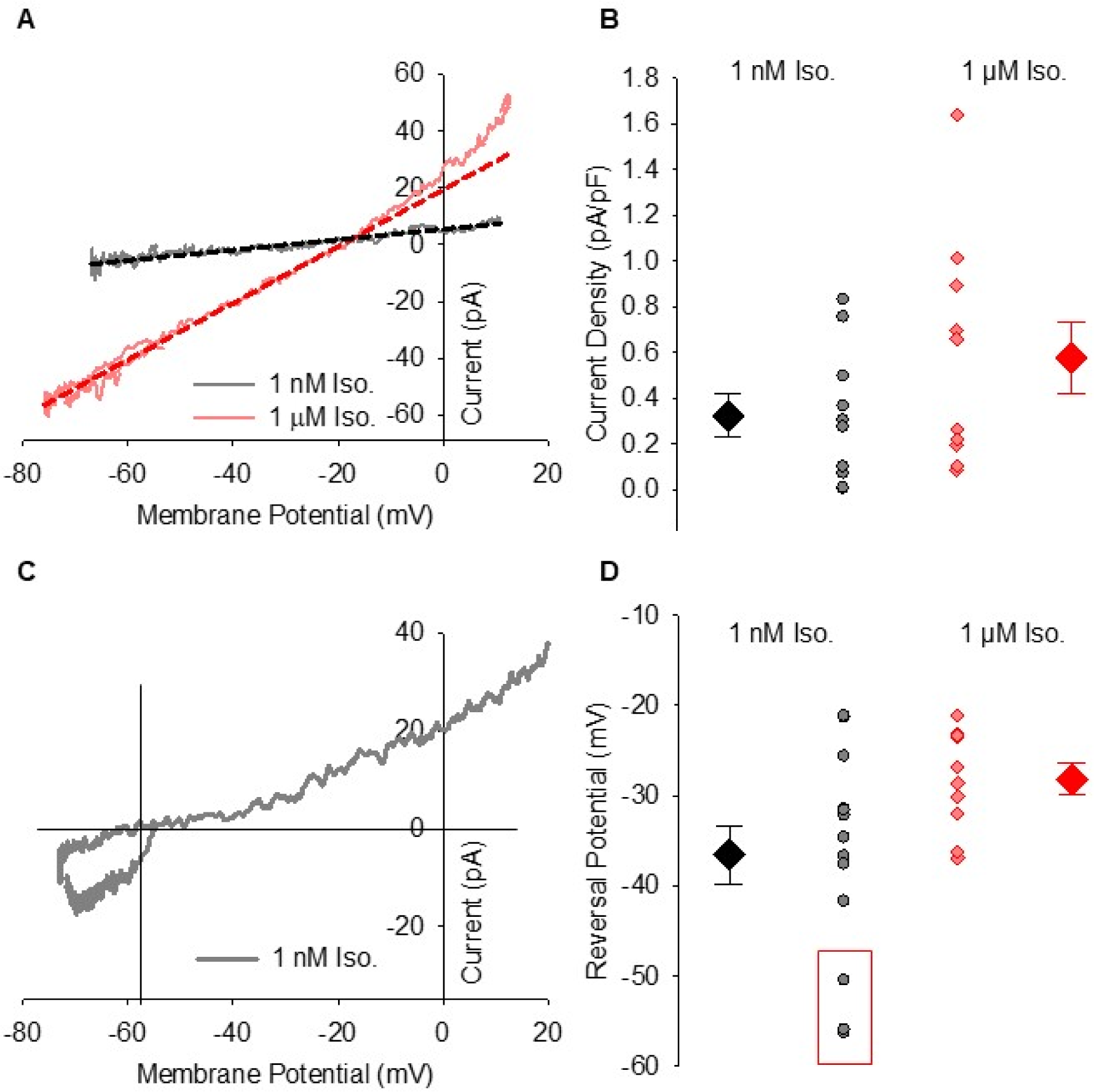
Rectification of the ivabradine-sensitive outward current at depolarized potentials. **A:** Representative experimental ivabradine-sensitive current-voltage relationships for a single cell in 1 nM isoproterenol (*black*) or 1 µM isoproterenol (*red*) with extrapolated linear fits to data at potentials negative to -20 mV indicated by *dashed lines*. **B:** Average (± SEM) peak outward I_f_ in SAMs in 1 nM isoproterenol (*black*) or 1 µM isoproterenol (*red*) predicted by extrapolated linear fits to experimental data at potentials below -20 mV. Individual data points are shown as small symbols. **C:** Representative ivabradine-sensitive current-voltage relationships from a cell with contamination by an outward current at diastolic potentials in 1 nM isoproterenol (*grey*). The reversal potential is -56 mV and there is apparent I_f_ activation during the diastolic depolarization as the contaminating current that opposes the inward I_f_ deactivates. **D:** Average (± SEM) reversal potential for ivabradine-sensitive current in 1 nM and 1 μM isoproterenol using the same colour scheme as **B**. The 3 cells included in the averages shown, but not included in the analysis in the main text are indicated by a red box. When these 3 cells are included the reversal potential of I_f_ differs significantly between 1 nM and 1 µM isoproterenol (unpaired t-test; P = 0.0494).

**Figure S4:**
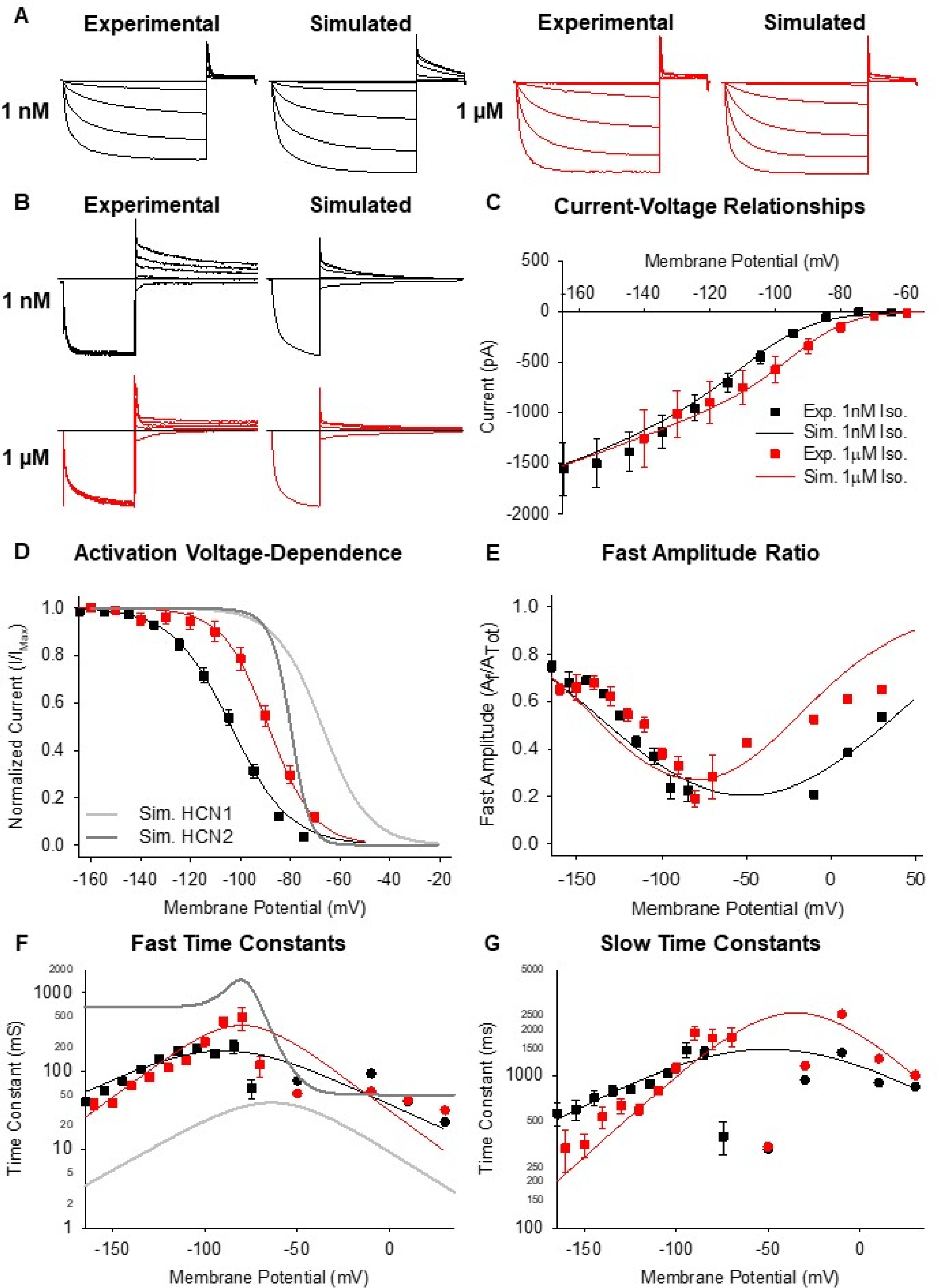
A model of the funny current recapitulates voltage-clamp data from SAMs. **A:** Experimental and simulated I_f_ current families in response to 3 s hyperpolarizations to potentials between -50 and -150 mV followed by a depolarizing step to +30 mV in 1 nM (*black*) and 1 µM (*red*) isoproterenol. **B:** Experimental (*left*) and simulated (*right*) I_f_ current deactivation in response to 3 s depolarizations to potentials between -50 and 30 mV following 1 s hyperpolarizations to -150 mV in 1 nM (*black*) and 1 µM (*red*) isoproterenol. **C:** Experimental (*symbols*) and simulated (*lines*) I_f_ current-voltage relationships in 1 nM (*black*) or 1 µM (*red*) isoproterenol. **D:** Experimental and simulated I_f_ activation voltage-dependence using the same colour scheme and symbols as **C**. Simulated HCN1 (*light grey*) (50) and HCN2 (*dark grey*) (51) activation voltage-dependence are also plotted. **E:** Experimental and simulated ratios of fast time constant to slow time constant amplitudes using the same colour scheme and symbols as **C**. **F-G:** Experimental and simulated fast (**F**) and slow (**G**) time constants of I_f_ activation (*squares*) and deactivation (*circles*) using the same colour scheme as **C**. Simulated HCN1 (*light grey*) and HCN2 (*dark grey*) time constants are also plotted in **F**. All experimental values are averages ± SEM. n = 5 to 20.

**Figure S5:**
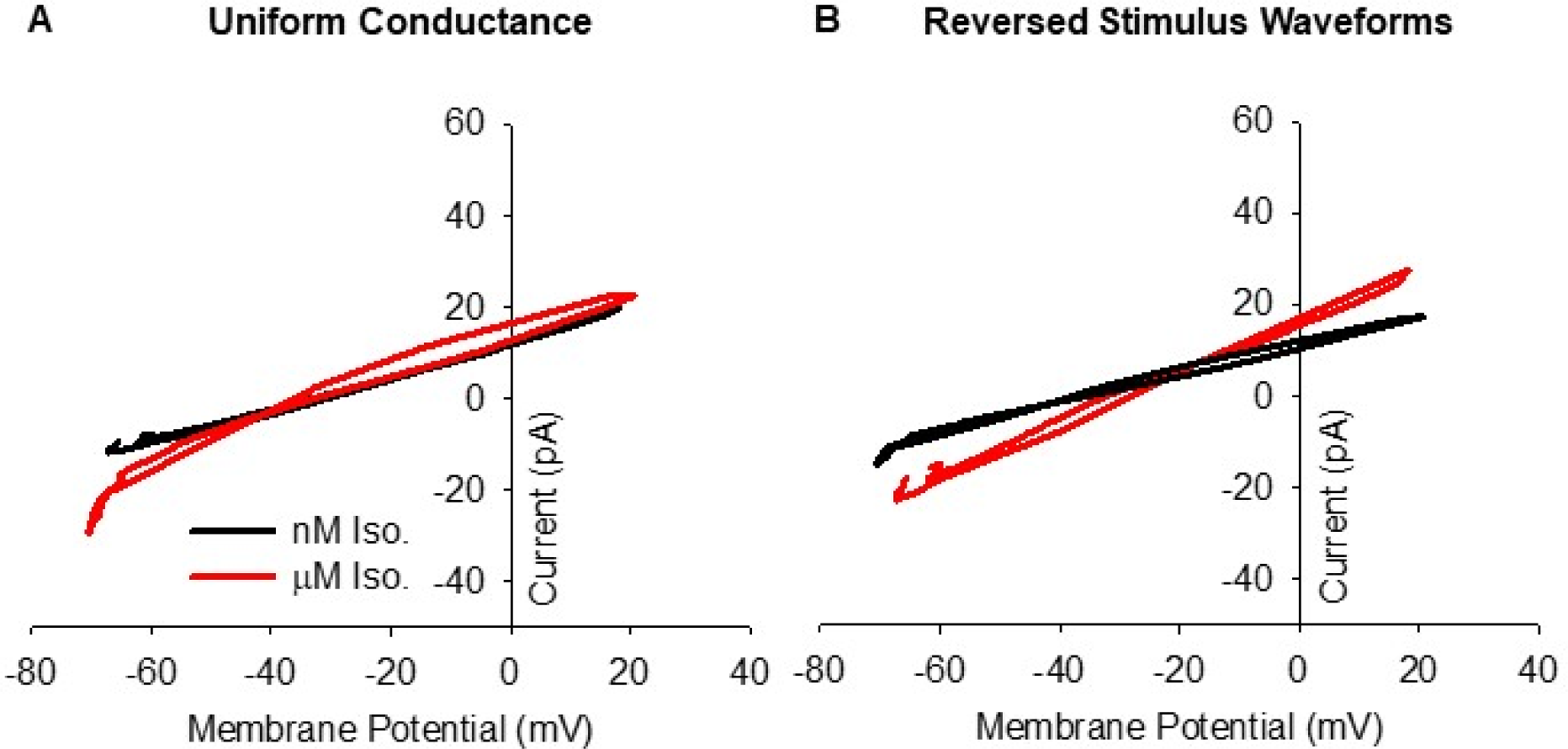
Changes in maximal conductance and AP waveform do not account for differences in the funny current in 1 nM and 1 µM isoproterenol. **A:** Average current-voltage relationships of simulated I_f_ in response to pre-recorded sinoatrial APs simulated using the 1 nM isoproterenol model (*black*) or the 1 µM isoproterenol model (*red*) with the same maximum conductance of 11.79 nS used in all simulations. **B:** Average current-voltage relationships of simulated I_f_ using the 1 nM isoproterenol model (*black*) simulated with APs recorded in 1 µM isoproterenol or the 1 µM isoproterenol model (*red*) with APs recorded in 1 nM isoproterenol.

## Footnotes

1 P-values, means, standard errors, and number of observations for all data are found in supplementary tables S1-S5.

## References

1. A. Keith, M. Flack, The Form and Nature of the Muscular Connections between the Primary Divisions of the Vertebrate Heart. J. Anat. Physiol. 41, 172–189 (1907).

2. D. DiFrancesco, The contribution of the “pacemaker” current (if) to generation of spontaneous activity in rabbit sino-atrial node myocytes. J. Physiol. 434, 23–40 (1991).

3. H. F. Brown, D. Difrancesco, S. J. Noble, How does adrenaline accelerate the heart? Nature 280, 235–236 (1979).

4. T. M. Ishii, M. Takano, L. H. Xie, A. Noma, H. Ohmori, Molecular characterization of the hyperpolarization-activated cation channel in rabbit heart sinoatrial node. J. Biol. Chem. 274, 12835–12839 (1999).

5. S. Moosmang, M. Biel, F. Hofmann, A. Ludwig, Differential distribution of four hyperpolarization-activated cation channels in mouse brain. Biol. Chem. 380, 975–980 (1999).

6. S. Moosmang, et al., Cellular expression and functional characterization of four hyperpolarization-activated pacemaker channels in cardiac and neuronal tissues. Eur. J. Biochem. 268, 1646–1652 (2001).

7. I. Ragueneau, et al., Pharmacokinetic-pharmacodynamic modeling of the effects of ivabradine, a direct sinus node inhibitor, on heart rate in healthy volunteers. Clin. Pharmacol. Ther. 64, 192–203 (1998).

8. A. J. Camm, C.-P. Lau, Electrophysiological effects of a single intravenous administration of ivabradine (S 16257) in adult patients with normal baseline electrophysiology. Drugs RD 4, 83–89 (2003).

9. A. Bucchi, A. Barbuti, D. Difrancesco, M. Baruscotti, Funny Current and Cardiac Rhythm: Insights from HCN Knockout and Transgenic Mouse Models. Front. Physiol. 3, 240 (2012).

10. D. DiFrancesco, Funny channel gene mutations associated with arrhythmias. J. Physiol. 591, 4117–4124 (2013).

11. A. O. Verkerk, R. Wilders, Pacemaker activity of the human sinoatrial node: an update on the effects of mutations in HCN4 on the hyperpolarization-activated current. Int. J. Mol. Sci. 16, 3071–3094 (2015).

12. E. G. Lakatta, D. DiFrancesco, JMCC Point-Counterpoint. J. Mol. Cell. Cardiol. 47, 157– 170 (2009).

13. V. A. Maltsev, E. G. Lakatta, Synergism of coupled subsarcolemmal Ca2+ clocks and sarcolemmal voltage clocks confers robust and flexible pacemaker function in a novel pacemaker cell model. Am. J. Physiol. - Heart Circ. Physiol. 296, H594–H615 (2009).

14. DiFrancesco Dario, The Role of the Funny Current in Pacemaker Activity. Circ. Res. 106, 434–446 (2010).

15. E. A. Accili, C. Proenza, M. Baruscotti, D. DiFrancesco, From Funny Current to HCN Channels: 20 Years of Excitation. Physiology 17, 32–37 (2002).

16. H. P. Larsson, How is the heart rate regulated in the sinoatrial node? Another piece to the puzzle. J. Gen. Physiol. 136, 237–241 (2010).

17. C. Altomare, et al., Integrated Allosteric Model of Voltage Gating of Hcn Channels. J. Gen. Physiol. 117, 519–532 (2001).

18. C. Proenza, D. Angoli, E. Agranovich, V. Macri, E. A. Accili, Pacemaker Channels Produce an Instantaneous Current. J. Biol. Chem. 277, 5101–5109 (2002).

19. C. Proenza, G. Yellen, Distinct populations of HCN pacemaker channels produce voltage-dependent and voltage-independent currents. J. Gen. Physiol. 127, 183–190 (2006).

20. A. O. Verkerk, R. Wilders, Hyperpolarization-activated current, If, in mathematical models of rabbit sinoatrial node pacemaker cells. BioMed Res. Int. 2013, 872454 (2013).

21. D. DiFrancesco, A. Ferroni, M. Mazzanti, C. Tromba, Properties of the hyperpolarizing-activated current (if) in cells isolated from the rabbit sino-atrial node. J. Physiol. 377, 61–88 (1986).

22. R. D. Nathan, Two electrophysiologically distinct types of cultured pacemaker cells from rabbit sinoatrial node. Am. J. Physiol. 250, H325–329 (1986).

23. M. R. Boyett, H. Honjo, I. Kodama, The sinoatrial node, a heterogeneous pacemaker structure. Cardiovasc. Res. 47, 658–687 (2000).

24. E. D. Larson, J. R. S. Clair, W. A. Sumner, R. A. Bannister, C. Proenza, Depressed pacemaker activity of sinoatrial node myocytes contributes to the age-dependent decline in maximum heart rate. Proc. Natl. Acad. Sci. 110, 18011–18016 (2013).

25. C. H. Peters, E. J. Sharpe, C. Proenza, Cardiac Pacemaker Activity and Aging. Annu. Rev. Physiol. 82, 21–43 (2020).

26. N. Haechl, J. Ebner, K. Hilber, H. Todt, X. Koenig, Pharmacological Profile of the Bradycardic Agent Ivabradine on Human Cardiac Ion Channels. Cell. Physiol. Biochem. Int. J. Exp. Cell. Physiol. Biochem. Pharmacol. 53, 36–48 (2019).

27. X. Wu, et al., Is ZD7288 a selective blocker of hyperpolarization-activated cyclic nucleotide-gated channel currents? Channels 6, 438–442 (2012).

28. C. F. Meier, B. G. Katzung, Cesium blockade of delayed outward currents and electrically induced pacemaker activity in mammalian ventricular myocardium. J. Gen. Physiol. 77, 531–547 (1981).

29. C. J. Abrams, N. W. Davies, P. A. Shelton, P. R. Stanfield, The role of a single aspartate residue in ionic selectivity and block of a murine inward rectifier K+ channel Kir2.1. J. Physiol. 493 (Pt 3), 643–649 (1996).

30. S. Zhang, S. J. Kehl, D. Fedida, Modulation of human ether-à-go-go-related K+ (HERG) channel inactivation by Cs+ and K+. J. Physiol. 548, 691–702 (2003).

31. Valenzuela Carmen, et al., Class III Antiarrhythmic Effects of Zatebradine. Circulation 94, 562–570 (1996).

32. M. Baruscotti, A. Barbuti, A. Bucchi, The cardiac pacemaker current. J. Mol. Cell. Cardiol. 48, 55–64 (2010).

33. J. L. Sánchez-Alonso, J. V. Halliwell, A. Colino, ZD 7288 inhibits T-type calcium current in rat hippocampal pyramidal cells. Neurosci. Lett. 439, 275–280 (2008).

34. J. P. Lees-Miller, et al., Ivabradine prolongs phase 3 of cardiac repolarization and blocks the hERG1 (KCNH_2_) current over a concentration-range overlapping with that required to block HCN4. J. Mol. Cell. Cardiol. 85, 71–78 (2015).

35. D. Melgari, et al., hERG potassium channel blockade by the HCN channel inhibitor bradycardic agent ivabradine. J. Am. Heart Assoc. 4 (2015).

36. P. Bois, J. Bescond, B. Renaudon, J. Lenfant, Mode of action of bradycardic agent, S 16257, on ionic currents of rabbit sinoatrial node cells. Br. J. Pharmacol. 118, 1051–1057 (1996).

37. C. Thollon, et al., Electrophysiological effects of S 16257, a novel sino-atrial node modulator, on rabbit and guinea-pig cardiac preparations: comparison with UL-FS 49. Br. J. Pharmacol. 112, 37–42 (1994).

38. A. Bucchi, M. Baruscotti, D. DiFrancesco, Current-dependent Block of Rabbit Sino-Atrial Node I_f_ Channels by Ivabradine. J. Gen. Physiol. 120, 1–13 (2002).

39. J. Stieber, K. Wieland, G. Stöckl, A. Ludwig, F. Hofmann, Bradycardic and Proarrhythmic Properties of Sinus Node Inhibitors. Mol. Pharmacol. 69, 1328–1337 (2006).

40. A. Bucchi, et al., Identification of the Molecular Site of Ivabradine Binding to HCN4 Channels. PLOS ONE 8, e53132 (2013).

41. M. E. Mangoni, et al., Functional role of L-type Cav1.3 Ca2+ channels in cardiac pacemaker activity. Proc. Natl. Acad. Sci. U. S. A. 100, 5543–5548 (2003).

42. M. Lei, et al., Requirement of neuronal- and cardiac-type sodium channels for murine sinoatrial node pacemaking. J. Physiol. 559, 835–848 (2004).

43. M. V. Brahmajothi, M. J. Morales, D. L. Campbell, C. Steenbergen, H. C. Strauss, Expression and distribution of voltage-gated ion channels in ferret sinoatrial node. Physiol. Genomics 42A, 131–140 (2010).

44. W. R. Giles, Y. Imaizumi, Comparison of potassium currents in rabbit atrial and ventricular cells. J. Physiol. 405, 123–145 (1988).

45. D. DiFrancesco, A study of the ionic nature of the pace-maker current in calf Purkinje fibres. J. Physiol. 314, 377–393 (1981).

46. D. DiFrancesco, Block and activation of the pace-maker channel in calf purkinje fibres: effects of potassium, caesium and rubidium. J. Physiol. 329, 485–507 (1982).

47. S. Kharche, J. Yu, M. Lei, H. Zhang, A mathematical model of action potentials of mouse sinoatrial node cells with molecular bases. Am. J. Physiol. Heart Circ. Physiol. 301, H945–963 (2011).

48. D. DiFrancesco, Characterization of single pacemaker channels in cardiac sino-atrial node cells. Nature 324, 470–473 (1986).

49. R. W. Tsien, Effects of Epinephrine on the Pacemaker Potassium Current of Cardiac Purkinje Fibers. J. Gen. Physiol. 64, 293–319 (1974).

50. A. Battefeld, N. Rocha, K. Stadler, A. U. Bräuer, U. Strauss, Distinct perinatal features of the hyperpolarization-activated non-selective cation current Ih in the rat cortical plate. Neural Develop. 7, 21 (2012).

51. J. Wang, S. Chen, M. F. Nolan, S. A. Siegelbaum, Activity-Dependent Regulation of HCN Pacemaker Channels by Cyclic AMP: Signaling through Dynamic Allosteric Coupling. Neuron 36, 451–461 (2002).

52. T. M. Ishii, M. Takano, H. Ohmori, Determinants of activation kinetics in mammalian hyperpolarization-activated cation channels. J. Physiol. 537, 93–100 (2001).

53. T. M. Ishii, N. Nakashima, K. Takatsuka, H. Ohmori, Peripheral N- and C-terminal domains determine deactivation kinetics of HCN channels. Biochem. Biophys. Res. Commun. 359, 592–598 (2007).

54. J. Stieber, et al., Molecular basis for the different activation kinetics of the pacemaker channels HCN2 and HCN4. J. Biol. Chem. 278, 33672–33680 (2003).

55. C. Altomare, et al., Heteromeric HCN1–HCN4 Channels: A Comparison with Native Pacemaker Channels from the Rabbit Sinoatrial Node. J. Physiol. 549, 347–359 (2003).

56. A. Ludwig, et al., Two pacemaker channels from human heart with profoundly different activation kinetics. EMBO J. 18, 2323–2329 (1999).

57. R. A. Capel, D. A. Terrar, The importance of Ca2+-dependent mechanisms for the initiation of the heartbeat. Front. Physiol. 6 (2015).

58. M. E. Mangoni, J. Nargeot, Genesis and Regulation of the Heart Automaticity. Physiol. Rev. 88, 919–982 (2008).

59. W. R. Giles, Supraventricular pacemaker activity in the canine heart: contributions from HCN channels in control conditions and in a model of heart failure. Cardiovasc Res 66, 430–2 (2005).

60. A. O. Verkerk, A. C. G. van Ginneken, R. Wilders, Pacemaker activity of the human sinoatrial node: Role of the hyperpolarization-activated current, If. Int. J. Cardiol. 132, 318–336 (2009).

61. Z. Liao, D. Lockhead, E. D. Larson, C. Proenza, Phosphorylation and modulation of hyperpolarization-activated HCN4 channels by protein kinase A in the mouse sinoatrial node. J. Gen. Physiol. 136, 247–258 (2010).

62. J. R. St Clair, Z. Liao, E. D. Larson, C. Proenza, PKA-independent activation of I_f_ by cAMP in mouse sinoatrial myocytes. Channels 7, 318–321 (2013).

63. M. Baruscotti, et al., Deep bradycardia and heart block caused by inducible cardiac-specific knockout of the pacemaker channel gene Hcn4. Proc. Natl. Acad. Sci. U. S. A. 108, 1705– 1710 (2011).

64. J. Alig, et al., Control of heart rate by cAMP sensitivity of HCN channels. Proc. Natl. Acad. Sci. U. S. A. 106, 12189–12194 (2009).

65. S. Herrmann, J. Stieber, G. Stöckl, F. Hofmann, A. Ludwig, HCN4 provides a “depolarization reserve” and is not required for heart rate acceleration in mice. EMBO J. 26, 4423–4432 (2007).

66. E. Hoesl, et al., Tamoxifen-inducible gene deletion in the cardiac conduction system. J. Mol. Cell. Cardiol. 45, 62–69 (2008).

67. T.-L. Lu, T.-J. Lu, S.-N. Wu, Inhibitory Effective Perturbations of Cilobradine (DK-AH269), A Blocker of HCN Channels, on the Amplitude and Gating of Both Hyperpolarization-Activated Cation and Delayed-Rectifier Potassium Currents. Int. J. Mol. Sci. 21, 2416 (2020).

68. M. Manz, M. Reuter, G. Lauck, H. Omran, W. Jung, A single intravenous dose of ivabradine, a novel I(f) inhibitor, lowers heart rate but does not depress left ventricular function in patients with left ventricular dysfunction. Cardiology 100, 149–155 (2003).

69. A. G. Torrente, et al., Burst pacemaker activity of the sinoatrial node in sodium–calcium exchanger knockout mice. Proc. Natl. Acad. Sci. 112, 9769–9774 (2015).

70. L. Dini, et al., Selective Blockade of HCN1/HCN2 Channels as a Potential Pharmacological Strategy Against Pain. Front. Pharmacol. 9 (2018).

71. M. Baruscotti, et al., A gain-of-function mutation in the cardiac pacemaker HCN4 channel increasing cAMP sensitivity is associated with familial Inappropriate Sinus Tachycardia. Eur. Heart J. 38, 280–288 (2017).

72. P. W. Liu, N. T. Blair, B. P. Bean, Action Potential Broadening in Capsaicin-Sensitive DRG Neurons from Frequency-Dependent Reduction of Kv3 Current. J. Neurosci. 37, 9705–9714 (2017).

73. T. Banyasz, B. Horvath, Z. Jian, L. T. Izu, Y. Chen-Izu, Sequential dissection of multiple ionic currents in single cardiac myocytes under action potential-clamp. J. Mol. Cell. Cardiol. 50, 578–581 (2011).

74. C. Rickert, C. Proenza, Action Potential Heterogeneity in Murine Sinoatrial Node Myocytes. Biophys. J. 112, 35a (2017).

75. Y. Chen-Izu, L. T. Izu, B. Hegyi, T. Bányász, “Recording of Ionic Currents Under Physiological Conditions: Action Potential-Clamp and ‘Onion-Peeling’ Techniques” in Modern Tools of Biophysics, Handbook of Modern Biophysics., T. Jue, Ed. (Springer, 2017), pp. 31–48.

76. E. J. Sharpe, J. R. S. Clair, C. Proenza, Methods for the Isolation, Culture, and Functional Characterization of Sinoatrial Node Myocytes from Adult Mice. JoVE J. Vis. Exp., e54555 (2016).

77. C. Rickert, C. Proenza, ParamAP: Standardized Parameterization of Sinoatrial Node Myocyte Action Potentials. Biophys. J. 113, 765–769 (2017).

